# Binary Brains: How Excitable Dynamics Simplify Neural Connectomes

**DOI:** 10.1101/2024.06.23.600265

**Authors:** Arnaud Messé, Marc-Thorsten Hütt, Claus C. Hilgetag

## Abstract

Fiber networks connecting different brain regions are the structural foundation of brain dynamics and function. Recent studies have provided detailed characterizations of neural connectomes with weighted connections. However, the topological analysis of weighted networks still has conceptual and practical challenges. Consequently, many investigations of neural networks are performed on binarized networks, and the functional impact of unweighted versus weighted networks is unclear. Here we show, for the widespread case of excitable dynamics, that the excitation patterns observed in weighted and unweighted networks are nearly identical, if an appropriate network threshold is selected. We generalize this observation to different excitable models, and formally predict the network threshold from the intrinsic model features. The network-binarizing capacity of excitable dynamics suggests that neural activity patterns may primarily depend on the strongest structural connections. Our findings have practical advantages in terms of the computational cost of representing and analyzing complex networks. There are also fundamental implications for the computational simulation of connectivity-based brain dynamics and the computational function of diverse other systems governed by excitable dynamics such as artificial neural networks.

---

The computational logic of most complex systems is based on the generation of local events and their global communication across the characteristic network architecture. The production of events is commonly based on the accumulation of inputs at susceptible nodes, which become excited when the inputs exceed a critical threshold value. Following their activation, nodes may enter a refractory period during which they cannot produce new events, before returning to a susceptible state. This basic cycle of activation and regeneration is the hallmark of excitable media. Despite the seemingly simple setup, this basic mechanism can generate rich global dynamics when combined with a heterogenous network organization [49].

Excitable dynamics can be found across a wide range of domains, from socio-economical and technological networks to biological systems [30, 102]. For example, communication between individuals in society is largely dictated by the exchange of messages, nowadays through social media [60, 67]. Biological excitable networks comprise diverse systems across multiple spatio-temporal scales, including interactions among species, cells or genes [66, 11]. A prime example of excitable systems are neuronal networks, composed of large populations of interconnected neurons, where the elementary signals are propagating action potentials [52].

Neuronal networks generate rich dynamics which are largely dictated by the underlying structural connectivity across scales [74, 93]. Thanks to technological progress, data on structural connectivity networks have been continuously refined. One major advance is the characterization of the strength of connections between neuronal entities [75, 59, 87], where broad skewed (lognormal) weight distributions have been observed across many scales of network organization [16]. For some aspects of systemic function, weights can serve as an important additional source of information and may even be essential for a mechanistic understanding of the system (see, e.g., [55]). To capture the functional implications of such weighted networks, suitable analysis techniques have been created and existing measures developed for binary graphs have been expanded [72, 7, 8, 83]. However, analyzing weighted networks presents several challenges due to the rich information they contain, which often requires a combination of advanced mathematical techniques and algorithmic innovation, making it a vibrant and evolving field on ongoing discussion and evolution. For example, there is as yet no universal definition of clustering in weighted graphs [2, 100], likewise how weights are transformed to topological distance remains an open question [4], see also [84]. As a conservative fallback, many analyses of the structural organisation of brain networks and the modeling of implications for brain activity and function are still being performed on binary rather than weighted versions of such networks. Given the attention on edge weights and their potential significance in network neuroscience, the limitations of unweighted graphs obtained, for instance, by imposing a binarization threshold have been widely debated [15, 89, 78, 96, 23, 105, 103, 81, 19]. At the same time, investigations of unweighted graphs have been widespread and very successful in revealing fundamental principles of biological neural networks [91, 9, 62], as well as networks in other contexts [38]. Consequently, the fundamental role played by network connection weights is still unclear, specifically, in terms of network communication.

How do the weights of a network influence the resulting excitation patterns? At the microscopic scale, it has been observed that spike propagation occurs mostly along strong and sparse links [73, 86, 37, 79], while at a larger scale, given the tight link between geometry and topology, it is likely that communication occurs mostly along proximal strong connections [39, 88]. Consequently, we hypothesize that the threshold encoded locally within an excitable system can be translated into a global network threshold, thereby unifying excitation patterns arising from weighted and binarized networks. By expressing the model excitation threshold as a function of the edge weights and assuming sparse excitation propagation, we show a direct mapping between dynamical patterns extracted from weighted networks and the ones extracted from binarized versions of the networks. We demonstrate this network-binarizing capacity of excitable dynamics in a stylized cellular automaton model and extend the framework to the FitzHugh-Nagumo model. Subsequently, we highlight the computational advantages of working with sparse binary graphs. We also apply the formalism to empirical brain connectivity data and find that simulations from binarized networks well reproduced dynamical features that have been empirically observed, including functional connectivity patterns as well as critical behavior. Finally, we illustrate the use of the formalism in the context of artificial neural networks, showing nearly equivalent accuracy of binarized networks compared to the weighted ones, accompanied by a drastic drop in models complexity. Thus, we find that, in a very generic fashion, excitable systems have the capacity to dynamically execute a threshold on the network weights, which has consequences for the functional interpretation of weighted networks.

## Main

### Overview

A weighted structural network can be represented as a graph where connections between nodes are summarized in a matrix *W*, whose elements *w_ij_* ≠ 0, if nodes *i* and *j* are connected, 0 otherwise. Connectome-based descriptions of neural dynamics are founded on a dynamical model describing the time evolution of the excitation patterns according to the local properties of the nodes, namely the activation function, and the global interactions among the nodes through *W*. By relating the intrinsic excitation threshold at the local node level (***‘model threshold’***) to the connection weights of the graph, we demonstrate a close correspondence between the activity patterns as observed in weighted networks to the ones of the associated binarized network versions (subject to a ***‘network threshold’*** at the global level). The primary dynamical feature that we investigate is the coactivation pattern, that is, the occurrence of simultaneous excitations between pairs of nodes, as a measure of functional connectivity (FC). In order to verify our predictions, we compare the patterns of coactivation across the two main parameters, the model threshold in weighted networks (dependent on the model parameters) and the network threshold applied to the connection weights. Subsequently, we extract the coactivation pattern across network thresholds that best match the coactivation pattern from the weighted network (***‘Matching’***), and compare the predicted threshold according to the maximal correlation against the model threshold (***‘Threshold agreement’***). See Fig 1 for the conceptual framework of the study. We then show that such binarizing capacity has computational advantages both in terms of memory footprint and execution time. Moreover, we apply the theoretical framework to empirical brain networks as well as artificial neural networks and find that, at the point where model threshold and network threshold are matched to each other, the simulations match well the empirical brain activity data, and artificial binary networks perform nearly as well as the weighted ones.

**Figure 1.**
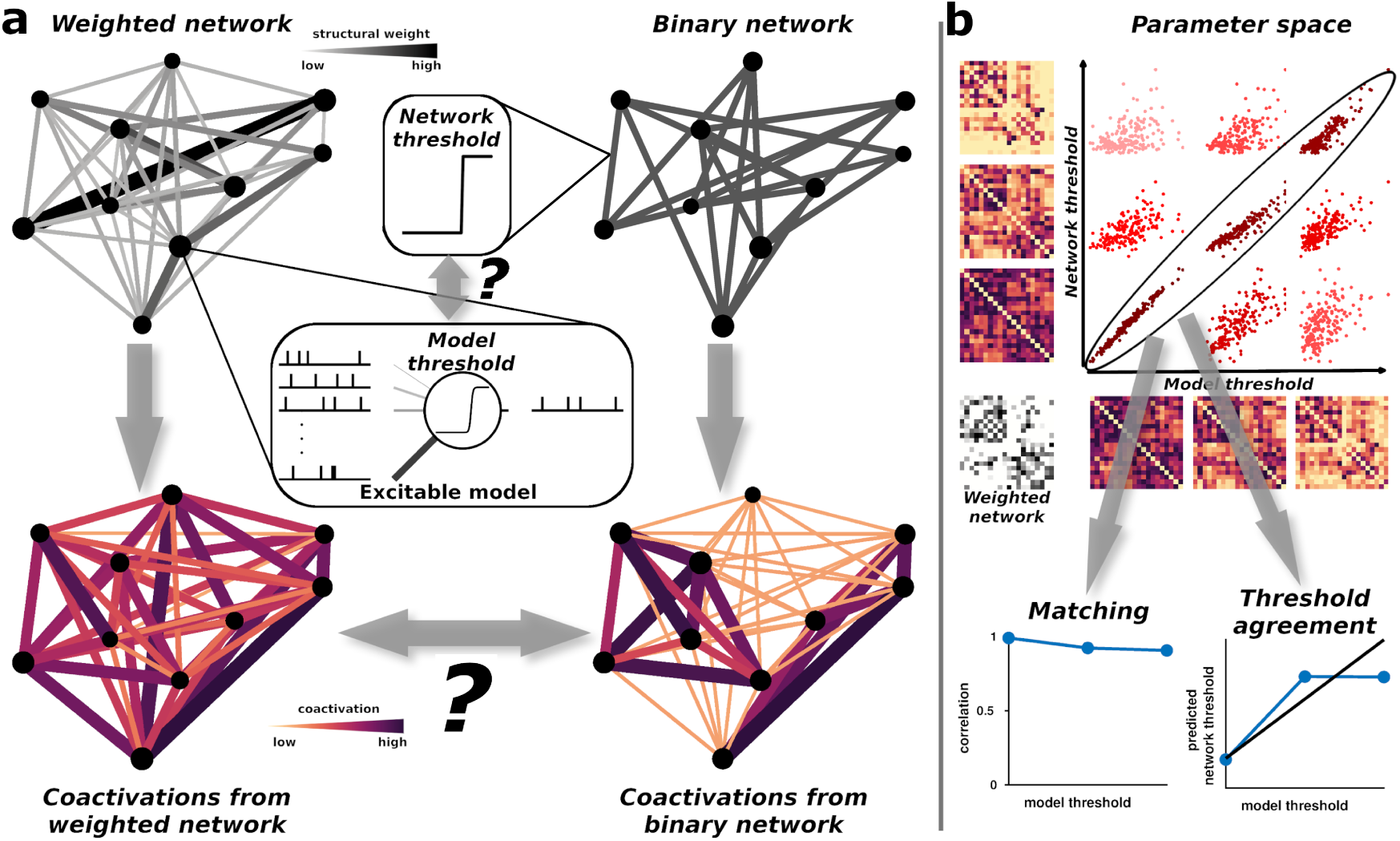
Illustration of the mapping of the model threshold to the network threshold. (a) Starting from a weighted network on which excitable dynamics run, we investigated how the local model threshold can be translated into a global network threshold. In an excitable network, fundamentally, a node’s activity is influenced by the inputs from the remainder of the network according to an activation function which encodes the model threshold. To verify the consistency of model versus network thresholds, we compared the resulting patterns of coactivations, a measure of functional connectivity. (b) Parameter space exploration of the association (correlation) between coactivation patterns from a weighted network with varying model threshold values and the coactivation patterns from binarized versions of the network with different network threshold values. Scatter plots represent coactivation from binarized networks versus coactivation from weighted networks. The color encodes the magnitude of the correlation coefficient. From this parameter space, we extracted the coactivation pattern across network thresholds that best matched the coactivation pattern from the weighted network (‘Matching’), and we compared the predicted threshold according to the maximal correlation against the model threshold (‘Threshold agreement’).

### Minimal excitable cellular automata model

As we strive for a mechanistic understanding of how weights influence dynamics, we initially study this question in a minimal model of discrete excitable dynamics, the SER model, following the Greenberg-Hastings formalism [42]. This minimal excitable model has a rich history in many disciplines, ranging from the propagation of forest-fires [5], the spread of epidemics [41], to neuronal dynamics [35]. Using such a model, we have previously shown, for example, that the distribution of excitations is regulated by the connectivity as well as by the rate of spontaneous excitations [69]. An increase in each of these two quantities leads to a sudden increase in the excitation density accompanied by a drastic change in the distribution pattern from a collective, synchronous firing to more local, long-lasting propagating excitation patterns. In a recent series of investigations, we have shown that the relationship between the network structure and the resulting coactivation patterns strongly and non-trivially depends on network topology [36, 49, 63, 62]. Qualitatively speaking, modularity is associated with a large positive structure-function correlation, while a broad degree distribution, or low network density, may lead to negative correlations.

In the SER model (Fig 2a), each node is assigned one of the three states: susceptible (S), excited (E) or refractory (R), and the transition rules are as follow: S → E with probability *f*, or if the sum of the connection weights with the excited neighbors is higher than a given model threshold *T*, i.e., Σ_*N_E_*_ *w > T* where N*_E_* being the number of excited neighbors of a node; E → R always; and R → S with probability *p*. In this setting, the activation function of the model is explicit and corresponds to a Heaviside step function shifted at *T*. We held fixed parameters *f* and *p*, which determine the time scales of self-excitation and recovery, respectively. To obtain insights into how the network weights are translated into the SER dynamics, we resorted to tools from mean-field theory. The general spirit of mean-field approaches is a space-implicit description of the system dynamics in terms of states probability (or density). Such approaches provide an average view of the dynamical space, by ignoring dependencies between nodes and inhomogeneities, that means, specific nodal connection patterns.

Let us first assume, for simplicity, a susceptible node in any binary graph with *k* neighbors (or degree). Using the SER model, with a model threshold value *T*, the node becomes activated, if more than *T* of its neighbors are excited. The corresponding probability, *P* (*X > T*), where *X* is the number of excited neighbors, is given by the complement of the cumulative binomial distribution

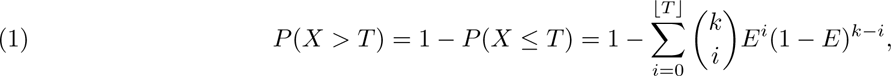

**Figure 2.**
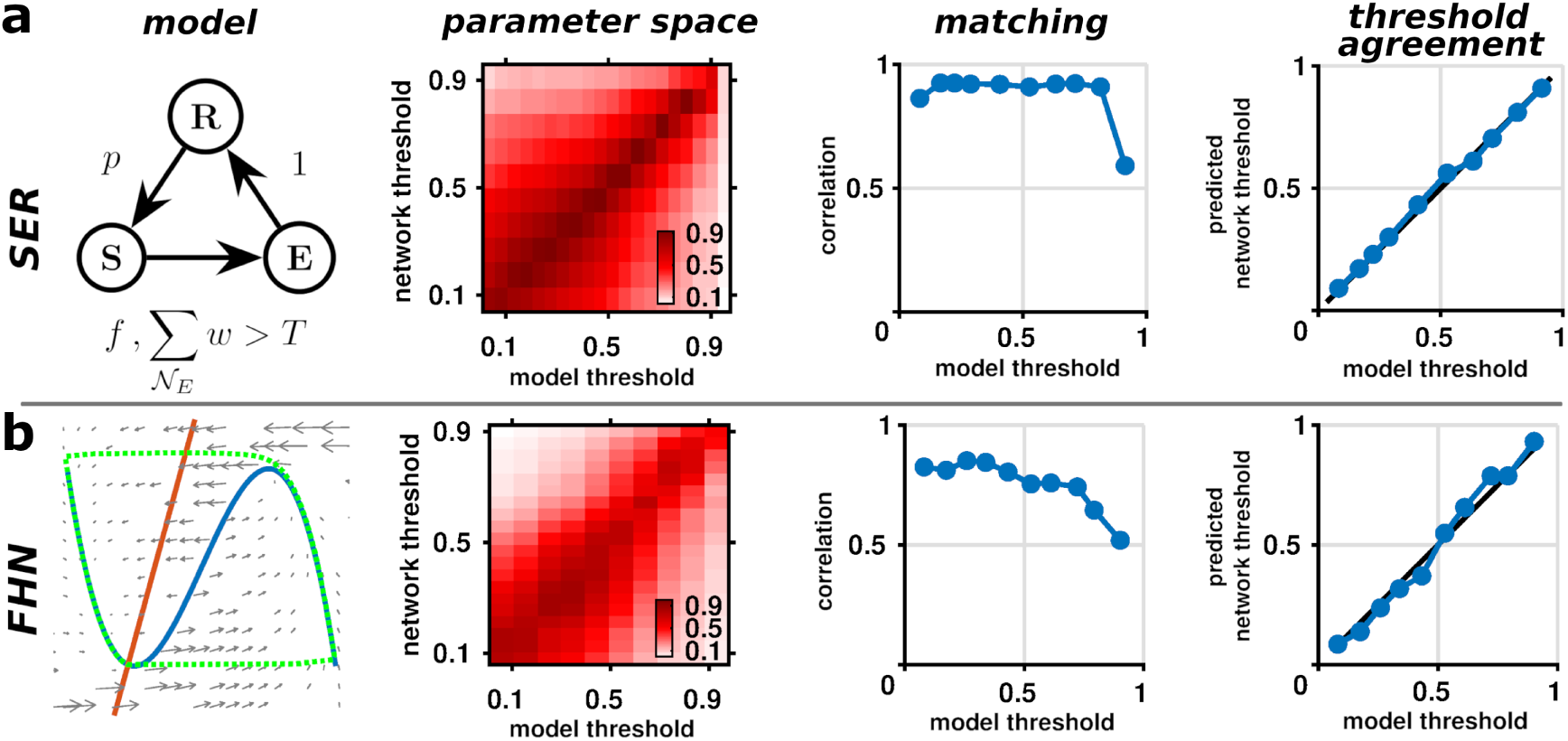
Summary of the main results. From left to right, schematic of the excitable models, parameter space exploration of the association between coactivation patterns from a weighted network with varying model threshold values and the coactivation patterns from binarized versions of the network with different network threshold values, the best match correlation (matching), and the corresponding predicted network threshold as a function of the model threshold (threshold agreement). The examples come from random weighted graphs with uniform weight distribution (N=100, k=4). For the SER model (a), *T*, N*_E_*, *p* and *f* represent the model threshold, the number of excited neighbors of a node, the recovery probability, and the spontaneous excitation probability, respectively. For the FHN model (b), phase plane, where blue and red curves represent the nullclines and the dotted green curve an exemplar trajectory.

where *E* corresponds to the density of excitation or the probability of a node being excited, and ⌊*T* ⌋ denotes the largest integer smaller or equal to *T*. The sum runs over all possible combinations of having a given number of excited neighbors *i* up to ⌊*T* ⌋. In the case where *T* = 0, the probability reduces to *P* (*X >* 0) = 1 − (1 − *E*)*^k^*, meaning that any active neighbor will trigger an excitation. When considering a node in any weighted network with arbitrary weight distribution, *P* (*W*), the terms within the sum are weighted by the corresponding cumulative distribution of having a given number of connections *i* (linked to excited neighbors) where the sum of their weights is lower or equal to 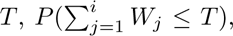 and then the probability that the node gets excited by its neighbors becomes

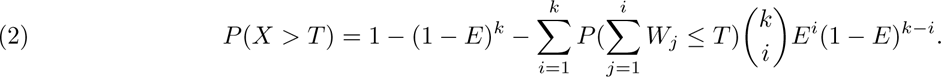

If we assume that the probability of having multiple neighbors excited simultaneously is negligible for a large range of *T* values (assuming implicitly that the excitation comes from a single neighbor), then we can neglect higher order terms and the probability reduces to

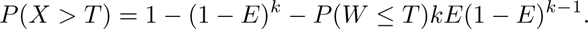

Now transforming the weighted network into a binarized version by applying a network threshold value *T_n_* equal to *T*, i.e., *P* (*W* ≤ *T_n_*) = 0, we observe that the model is equivalent to the one with *T* = 0. That is, running the SER model on a weighted graph with a model threshold value *T* is equivalent to running the model on a binarized version of the graph using a network threshold value *T_n_* = *T* and a model threshold *T* = 0. In that way, our hypothesis of a threshold mapping the dynamics on weighted graphs onto binarized graphs is demonstrated by such theoretical arguments. We numerically verified the existence of this threshold in an exemplar random network with uniform weight distribution where we observed a near perfect agreement (Fig 2a). We subsequently performed an extensive validation, see SI, showing that the mapping is valid across diverse network structures (including various network topologies and weight distributions). Interestingly, we observed that the best predictions are obtained from log-normal distributions, ubiquitous in empirical data. As such, assuming that excitations are coming from single neighbors appears to be valid, this assumption is verified in detail in SI Number of excited neighbors.

### Fitzhugh-Nagumo model

Second, we studied the FHN model as an established low-dimensional system of excitable dynamics [32, 70]. The FHN model is a two-dimensional simplification of the classical biophysical neuron model of Hodgkin and Huxley [47], composed of two nested variables, the membrane potential *u* and a recovery variable *v*

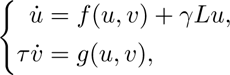

where 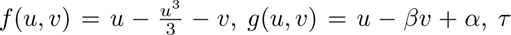 represents the time scale of the recovery variable fixed to 100, and *γ* is a global scaling parameter or coupling strength modulating the network influence. By introducing the network Laplacian matrix, *L* = *W* − *D*, where *D* is the strength (or degree for binary networks) matrix, *D_ij_* = Σ*_k_ w_ik_* if *i* = *j*, zero otherwise, the diffusive coupling between elements of the network is expressed as *γLu*.

Depending on the values of the governing parameters (*α* and *β*), each node may be in an excitable, oscillatory or non-excitable steady state regime. The phase plane of the FHN model allows a geometrical explanation of the dynamical regimes related to neuronal excitability, i.e., the presence of a threshold and spike-generating mechanism (Fig 2b). This phase plane is characterized by the nullclines of the system, namely when *f* (*u, v*) = *g*(*u, v*) = 0. The intersection of the nullclines represents the fixed-points (*u*^∗^*, v*^∗^). We are here interested in the excitable regime of the model which corresponds to having only one stable fixed point. By fixing *β* = 0.6, the excitable regime is established for *α >* 0.6. As opposed to the SER model, the FHN model does not have an explicit activation function, and hence a trivial well-defined threshold, due to the absence of a saddle equilibrium [31]. We thus need first to extract the intrinsic model threshold from the phase plane caracteristics.

Let us first assume a brief pulse input (of width *dt*) to a node. Intuitively, for a spike to occur, the magnitude of the input *I* must be greater than the distance, denoted *d*, separating the fixed point from the unstable branch of the N-shaped nullcline, *I > d/dt* (see SI FHN model threshold)). When considering a network, we suppose that a spike is propagated to a neighbor if the energy transferred (E) multiplied by the network effect (i.e., the connection weight *w* times the coupling strength *γ*) allows the variable *u* to exceed the distance *d*. Consequently, we obtain the condition on network weights for spike propagation

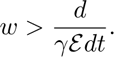

While the distance *d* can be easily computed analytically as a function of *α*, we do not have a clear and simple expression for the energy E transferred by a spike through the network. Therefore, we resorted to numerical experiments (see SI FHN model threshold)). Thus, using the same reasoning as for the SER model, we can potentially binarize the weighted graph by applying a network threshold that satisfies the previous condition to obtain similar dynamics and consequently similar coactivation patterns. Subsequently, the parameter *α* was fixed to 0.61. We verified our prediction in an exemplar random network with uniform weight distribution and observed a generally good agreement across a large range of parameter values (Fig 2b). As for the SER model, we subsequently performed an extensive validation, see SI, showing the validity of the mapping across diverse network structures (with varying topology and weight distribution), and stronger agreements on log-normal distributions.

### Computational consequences

In this section, we show that, beyond the theoretical aspects of the network-binarizing capacity of excitable dynamics, thresholded and binarized networks present computational advantages both in terms of memory and execution time. To determine these computational aspects, we used random weighted networks with an uniform weight distribution and a fixed average degree of 20 and varying number of nodes (from 500 to 20,000). Subsequently, these networks were thresholded at different values (0.25, 0.5 and 0.75). All the experiments were performed using Matlab (version 9.12.0 (R2022a), Natick Massachusetts: The MathWorks Inc. https://www.mathworks.com) on an AMD EPYC 75F3 server with 32 CPU cores at 2.95 GHz with 1 TB RAM. Simulations were repeated 5 times and the median execution times are shown.

First, the memory requirement to represent binary networks is much lower than for weighted networks, and the gain compared to weighted networks increases with increasing network size by several order of magnitude. For example, a network with 20,000 nodes requires about 1 GB of memory for the weighted version, while thresholded and binarized versions only need about 1 MB (Fig 3a). Generally, thresholded and binarized networks have low connections density and can therefore be represented by sparse formats. Moreover, we can use logical data types to represent connections, which further reduces memory requirement. That is, it requires just one byte to store a connection while weights need at least 4 bytes. Second, we explored the execution time of the SER model (*p* = 0.1; *f* = 0.01; 10,000 time steps). Again, due to the sparsification of the networks following thresholding and binarization, execution time was also greatly reduced, from ∼10 min for a weighted network with 20,000 nodes to only few seconds for its binarized versions (Fig 3b).

**Figure 3.**
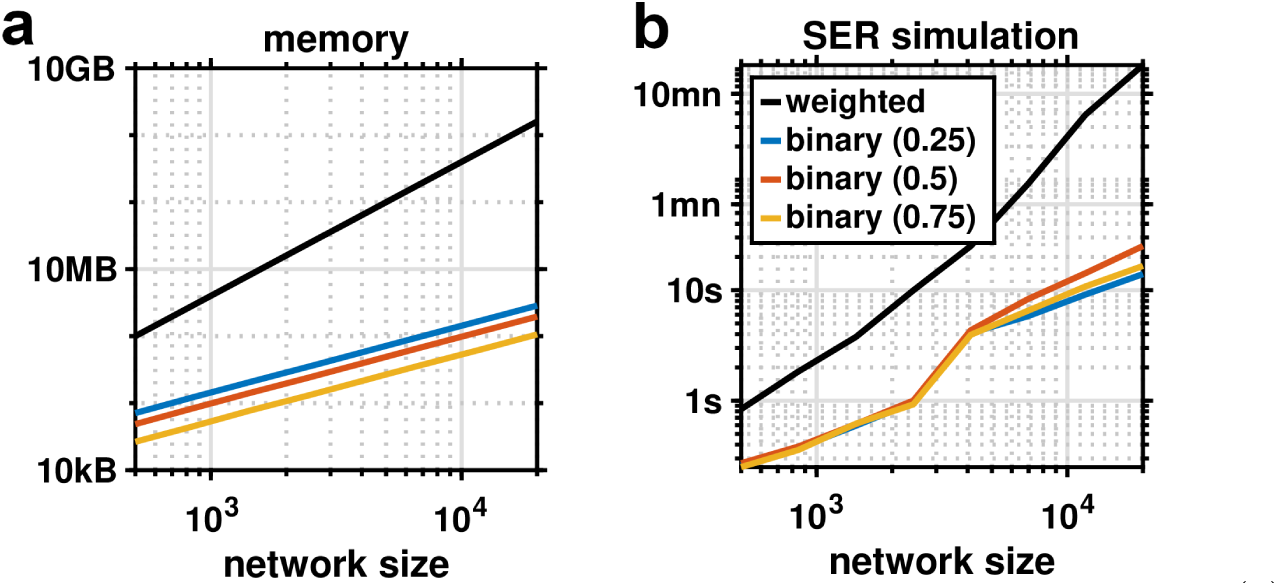
Computational aspects. Network memory footprint in bytes (a) and SER model execution time (b) as a function of the number of nodes (x-axis) and the network type (weighted or thresholded and binarized).

The computational advantages of binarized networks are not only observed in dynamical simulations, but also when computing basic matrix operations as well as topological caracteristics of the networks and this through diverse software environments, see SI Computational advantages.

### Empirical brain networks

Next, we applied the outlined theoretical framework using the SER model to an empirical dataset of structural and functional brain connectivity derived from MRI data of 70 healthy participants [43] (Fig 4a, see Materials and Methods section for details).

**Figure 4.**
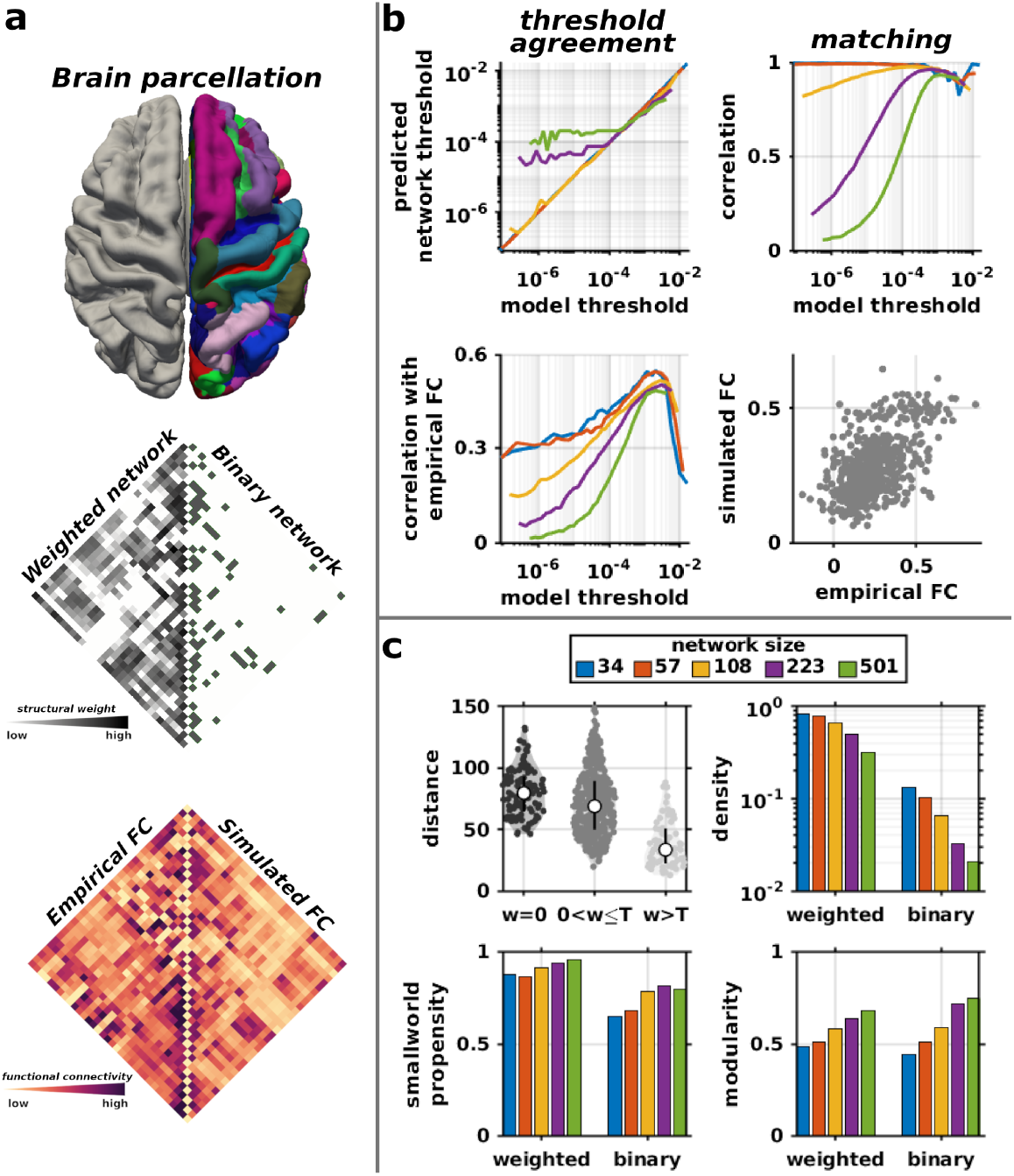
Exploration of empirical brain networks. (a) From top to bottom, the brain parcellation of the right hemisphere (at the coarsest scale of 34 regions), the original weighted structural network and the corresponding binarized version for which the correlation with the empirical FC was maximal, the empirical FC and the best match simulated FC. (b) Threshold agreement, matching and the correlation between simulated and empirical FC as a function of the model threshold (or equivalently the network threshold). Colors code for different network size. The corresponding scatter plot between the empirical FC and the best match simulated FC for the coarsest scale. (c) Distribution of the Euclidian distance (mm) between pairs of nodes according to the structural weight, where *T* denotes the threshold for which the correlation between the empirical and simulated FCs is maximal, for the coarsest scale. The density, smallworld propensity and modularity of the original weighted structural networks and the binarized versions. Colors code for different network parcellation sizes.

Similar parameter setting as for the synthetic networks were employed with the empirical brain networks when performing SER simulations. We extracted the coactivation pattern across network thresholds that best match the coactivation pattern from the weighted network, and compare the predicted threshold according to the maximal correlation against the model threshold. As for the synthetic networks, predictions generally worked well across spatial scales (considering brain network parcellations from 34 up to 501 regions) and threshold values. Only the predictions for large networks deviated from the theory (Fig 4b).

Additionally, in order to appreciate the ability of the excitable model to predict empirical functional connectivity, we computed the correlation between the empirical and simulated FC. For that purpose, simulated excitation patterns were first convolved with the standard hemodynamic response function to model the neurometabolic coupling, before computing simulated FC as the zero-lag Pearson correlation coefficient between all pairs of brain regions. We observed that the prediction depends on the model threshold (or equivalently on the network threshold). There was an optimal threshold value maximizing the correspondence of the simulations with the empirical data. Across spatial scales, the maximum correlation was close to 0.6, consistent with previously published models of empirical brain activity [92, 93].

Given the empirical relationship (i.e., anti-correlation) between the structural weights and Euclidian distance [80], the threshold value maximizing the association with the empirical FC appeared to remove mostly long-distance connections, where the distances were of the same order as those for unconnected regions pairs (Fig 4c). Of note, some long-range connections survived the threshold value. The resulting binarized networks were about one order of magnitude sparser than the original weighted networks. Interestingly, despite such changes in density, the binarized networks presented similar topological properties in terms of smallworldness and modularity as the original ones (Fig 4c).

Binarized networks also reproduced a number of dynamical properties reminescent of critical behavior, see SI Brain criticality.

### Artificial neural networks

Finally, we explored the possibility to transfer the current theoretical framework to the context of artificial neural networks (Fig 5). For that purpose, we resorted to the perceptron model, the first artificial neural network invented in 1958 by psychologist Frank Rosenblatt [82]. We implemented a simple model to classify handwritten digits using the classic benchmark image classification dataset MNIST [27], see Materials and Methods section for details.

**Figure 5.**
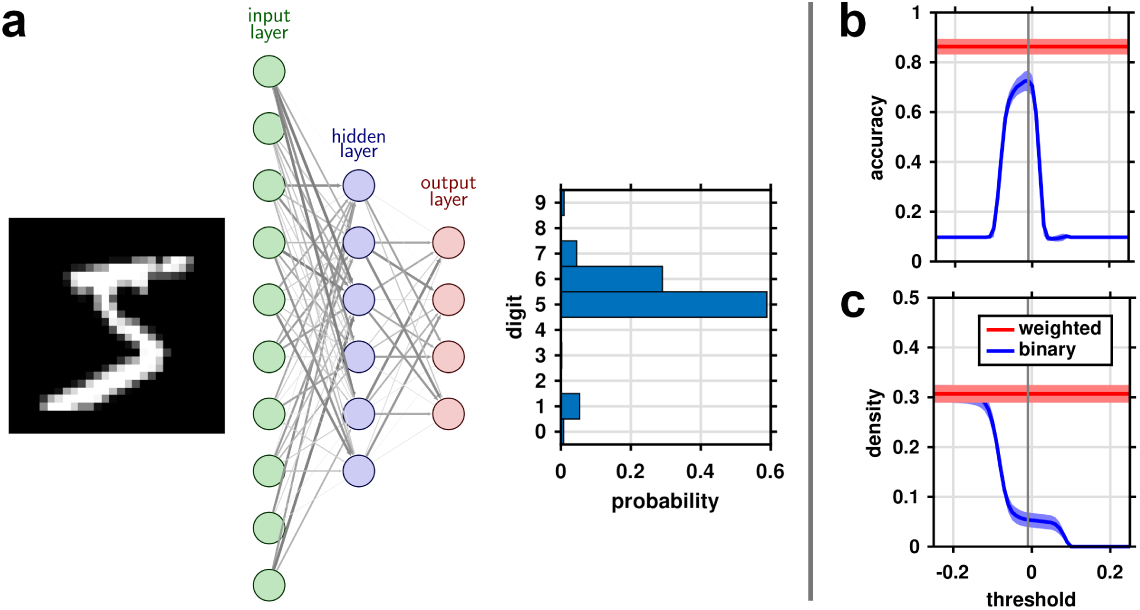
Exploration of artificial neural networks. (a) The perceptron model. Pixels of the input image are fed to the input layer units, processed through the hidden layer units, and finally output layer units code the probability that the image represents a given digit. (b) Accuracy as a function of the network threshold of the binarized networks (blue) and the corresponding accuracy of the original weighted networks (red). (c) Proportion of connections from the input layer to the hidden layer for the original weighted networks (red) and the thresholded and binarized networks (blue). Curves and shaded areas represent the average and standard deviation over 50 training realizations.

Once the model was trained, we thresholded and binarized the connections from the input layer to the hidden units at various threshold values and subsequently evaluated the performance of the resulting network. The performance of the binarized network approached the one from the original weighted network near the theoretical threshold, here set to zero (Fig 5b). Interestingly, the maximal accuracy was obtained for very sparse networks with only 5% of connections compared to the 30% for the weighted networks, on average (Fig 5c). The total number of parameters (including the number of connections and bias) dropped on average from about 76,222 to 15,070, i.e., 80% of the parameters were removed.

## Discussion

Understanding the mechanisms of communication in complex systems is a central focus of network science. Such an understanding not only allows to predict dynamical behaviors, but also, potentially, to manipulate and steer the system away from unwanted or pathological states [58]. Our present investigations point to a straightforward mapping between excitable dynamics implementing an intrinsic threshold running on complex weighted networks versus on binarized versions of these networks. Mapping the model threshold into network weights, we demonstrated a direct agreement between coactivation patterns based on weighted networks and the ones derived from binary versions of the same networks.

Due to recent technological advances, large-scale recordings of neuronal populations together with their synaptic connectivity have become feasible [98]. Empirical evidence of heavy tail synaptic distributions [16] has been corroborated with its functional importance, such as the presence of strong connections between neuronal populations that show correlated activity and similar responses [95, 104, 53, 24, 56]. A number of recent studies have started to highlight general principles of spike communication. Overall, these studies have demonstrated that spike transmission occurs mostly along strong and sparse links [73, 86, 37, 79]. At the macroscopic scale, communication models have shown that interactions occur mostly along proximal strong connections, a consequence of the tight link between geometry and topology [39, 88, 93]. Moreover, the heavy-tailed distributions of synaptic weights enhance the dynamical repertoire of the system, and increase its response speed and stability [51, 73, 86, 79]. However, given the predominance of strong links in communication, it remains unclear whether weak links are of functional importance or whether they can be discarded without noticeable consequences on the resulting dynamics.

In the present study, we specifically tested and subsequently supported the importance of strong links by showing a clear correspondence between excitable dynamics running on complex weighted networks with a simple binarized version of these networks. This equivalence, from a dynamical perspective, can explain the success of structural analyses performed on binary graphs, and the wide range of design principles this simplified perspective of complex systems has helped to elucidate. Moreover, our results showed that excitation propagates through networks via cascade events, i.e., excitations are triggered by a single neighbor, which appears to be consistent with the mounting empirical evidence of such mechanisms in brain dynamics [46, 57, 54]. The thresholding capability, as a generic feature of excitable dynamics, points to the need of a careful empirical investigation of other features, such as the precise value of the coupling strength, which, in turn, implements the thresholding capabilities and the resulting value of the effective threshold generated by the excitable model. Consequently, the coupling strength of excitable models may serve as a lens into the network topology, where the aperture variation allows to highlight the multiscale network organization [3, 106].

The results also echo the concept of effective connectivity, defined as the influence that a node exerts over another [34]. The mapping of the excitation threshold on network weights conveniently reveals the connections that are effectively used functionally. Further investigations are needed to probe such potential bridges between these two concepts. As an illustrative example, we explored our framework for empirical brain networks. We observed a model threshold, and a corresponding network threshold, for which the simulated FC reproduced the empirical FC pattern within the range observed by diverse computational models incorporating more biological realism [64, 17]. These results are consistent with a number of studies which demonstrated the ability of simple models to predict FC, and sometimes even outperform more complex ones [45, 39, 65, 61, 85]. The most intriguing result here is that such predictions can be, all the more, computed from a sparse binarized version of the structural networks. This threshold resulted in quite sparse and mostly short-range binary networks, but preserved the topological characteristics of the original weighted networks. Interestingly, a handful of studies have started to explore the effect of thresholding (but without binarization) in various contexts. It has been shown that, similar to our results, thresholding weak connections (up to ∼ 90%) has no effect on the topological properties of the structural networks [22]. Connectivity gradients, a low-dimensional representation of functional interactions, are robust against the full range of connectivity strength thresholds [71]. Furthermore, thresholding enhances network-age association, by potentially removing spurious connections [14].

Of note, while the threshold mostly cuts long-distance connections which typically have a low weight, we observed that some of them still survive. These connections are likely the ones deviating from the distance rule, i.e., they possess strength values higher than expected given their length [80, 10]. All the more, longrange brain connections play fundamental roles, for instance, by increasing dynamic complexity and linking high-order cognitive functions [64, 20, 10, 99], a role resembling Granovetter’s notion of the ‘strength of weak ties’ [40]. Our results show that, when it comes to predicting the pattern of functional connectivity, the predominant factor is the set of short (and strong) connections, demonstrating that such a scaffold of strong connections effectively constraints functional interactions. This finding also appears to be borne out by other recent work [76].

The present results also have implications for the design of artificial neural networks. Classically, neural networks adjust their connection weights during training in order to minimize prediction error. Moreover, typical artificial neuron models, such as the famous McCulloch-Pitts neuron, are reminiscent of the excitable model formalism explored in the present study. As an illustrative example, we explored our framework for the perceptron model, where we observed also a similar straightforward mapping between excitable dynamics from weighted versus binary networks together with nearly similar accuracy. Interestingly, binarized neural networks show promise as an emerging alternative in machine learning, providing similar performances but with spectacular reduction in computational burden, memory footprint and energy consumption [48, 90], observations that we also reported here in a more general context.

Importantly, our framework is void of strong constraints and can therefore generalize to various alternative scenarios (see SI Generalization). The threshold equivalence is valid for diverse network configurations, such as directed networks or when the weight distribution depends on the underlying topology [7]. It also generalizes to variations in the dynamical model, such as when network nodes have heterogeneous threshold values [50] or when the activation function is probabilistic.

As a potential limitation of the present approach, we find that the agreement between simulations on the weighted and unweighted graphs is predominantly determined by the distribution of required neighbors triggering an excitation (which in turn is influenced by the network’s weight distribution). If this distribution of required neighbors is narrow and centered around one neighbor, the agreement is high. By contrast, if this distribution is broad or centered around a larger number of neighbors, we cannot compute a suitable threshold translating the weighted graph into the unweighted graph.

In summary, we demonstrate how local thresholds imposed by excitable models translate into global thresholds on network weights, revealing a sparse scaffold of strong connections that are the most effective drivers of network dynamics.

## Materials and Methods

### Excitable models

#### SER model

We used a simple three-state cellular automaton model of excitable dynamics, the SER model. The SER model operates on discrete time and employs the following synchronous update rules, a node in the:

- S → E at probability *f* or if the sum of the weights connecting excited neighbors is greater than *T*;
- E → R;
- R → S at probability *p*.

For *f* = 0 and *p* = 1, one obtains the deterministic version which was investigated in detail in [36, 62], where, for example, the role of cycles in storing excitations and supporting self-sustained activity, and the generative mechanisms of coactivations were elucidated. Here we employed the model in a stochastic regime where *f >* 0, *p <* 1. We used fixed parameters *f* = 0.01 and *p* = 0.1. The only remaining parameters were the underlying topology and weight distribution of the connectivity network on which the model runs and the threshold value *T*.

To analyze the pattern of joint excitations, we computed the number of simultaneous excitations between all pairs of nodes. The outcome matrix is the so-called coactivation matrix, a measure of the functional connectivity of the network:

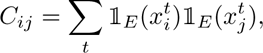

where 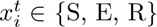 being the state of node *i* at time *t*, and 1*_E_* the indicator function of state E

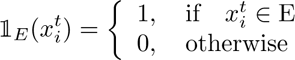

For each network realization and threshold value, we computed 50 runs of 10,000 time steps each. We increased the number of runs to 100 and the number of time steps to 100,000 for the simulations on empirical data. The initial conditions were generated randomly, with 10% of nodes excited while the remaining nodes were equipartitioned into susceptible and refractory states. The model threshold values were set as the decile of the corresponding weight distribution, while the network threshold values were set as the trentile of the associated weight distribution. The model threshold when performing simulations on binarized networks was set to zero.

#### FHN model

We used the popular Fitzhugh-Nagumo model [32, 70]. The FHN model is governed by the following differential equations:

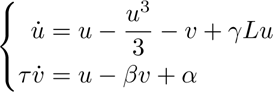

Unless otherwise stated, parameter values are set such that the model is in an excitable regime with *α* = 0.61 and *β* = 0.6.

To analyze the pattern of joint excitations we used a method developed earlier [63]. In particular, given the comparatively broad spikes in the FHN model compared to the SER model which leads to an imprecise separation of coactivations and sequential activations, time-series were discretized and spikes detected. The detection threshold for spikes was determined as zero, so any onset activity above this threshold was considered as spikes. Then, coactivation patterns were computed as for the SER model. A time window of 20 ms is used from which ‘simultaneous’ events were drawn for computing coactivations.

Simulations were performed using Euler integration with an assumed time resolution of 0.1 ms, and further subsampled at 1 ms. Each run lasted 100 s. In order to sustain the activity within the network, nodes were driven by spontaneous Poisson noise with a rate of 0.05 Hz. The coupling parameter value (*γ*) was set as the ratio between the model threshold and the decile of the corresponding weight distribution, while the network threshold values were set as the trentile of the associated weight distribution. The coupling value when performing simulations on binarized networks was set to the model threshold value.

### Empirical brain networks

To further verify our predictions, we also analyzed an independently collected and preprocessed dataset from the Lausanne University Hospital [43]. Informed consent was obtained for all subjects (the protocol was approved by the Ethics Committee of Clinical Research of the Faculty of Biology and Medicine, University of Lausanne, Switzerland, #82/14, #382/11, #26.4.2005). A total of 70 healthy participants (age 28.8 ± 9.1 years, 27 females) were scanned in a 3-Tesla MRI Scanner (Trio, Siemens Medical, Germany). Details regarding data acquisition, pre-processing and network reconstruction are available at [43].

Briefly, the data acquisition protocol included a magnetization-prepared rapid acquisition gradient echo (MPRAGE) sequence (1 mm in-plane resolution, 1.2 mm slice thickness), a diffusion spectrum imaging (DSI) sequence (128 diffusion-weighted volumes and a single b0 volume, maximum b-value 8,000 *s/mm*^2^, 2.2×2.2×3.0 mm voxel size), and a gradient echo-planar imaging (EPI) sequence sensitive to blood-oxygen-level-dependent (BOLD) contrast (3.3 mm in-plane resolution and slice thickness with a 0.3 mm gap, TR 1,920 ms, resulting in 280 images per participant). The Connectome Mapper Toolkit was used for the initial signal processing [26] while gray and white matter were segmented from the MPRAGE volume using Freesurfer [28]. Cortical grey matter was first parcellated into 68 areas using Freesurfer [28], and then further subdivided into 114, 219, 448 or 1,000 equally sized parcels [18].

Structural connectivity matrices were reconstructed for individual participants using deterministic streamline tractography on reconstructed DSI data. 32 streamline propagations were initiated per diffusion direction and per white matter voxel [101]. The weights of the edges correspond to the streamline density. fMRI volumes were corrected for physiological variables (regression of white matter, cerebrospinal fluid, as well as motion), BOLD time-series were subjected to a lowpass filter and motion “scrubbing” [77] was performed. Functional connectivity matrices were constructed by computing the zero-lag Pearson correlation coefficient between the fMRI BOLD time-series of each pairs of brain regions. Group-average structural and functional connectivity matrices were reconstructed simply by averaging all individual connectivity matrices. Given the intrinsic drawbacks of tractography algorithms, namely their difficulty to properly detect crossing fibers as well as to track long distance fibers [64, 97], we focused only on the intra-hemispheric connectivity of the right hemisphere.

Similar parameter setting as for the synthetic networks were employed with the empirical brain networks when performing SER simulations (i.e., *f* = 0.01 and *p* = 0.1). For each threshold value, we computed 100 runs of 100,000 time steps each. The model threshold values were set as the trentile of the empirical structural weight distribution, while the network threshold values were set as the 60-quantile of that distribution. Subsequently, we extract the coactivation pattern across network thresholds that best match the coactivation pattern from the weighted network, and compare the predicted threshold according to the maximal correlation against the model threshold.

Additionally, in order to appreciate the ability of the excitable model to predict empirical functional connectivity, we computed the correlation between the empirical and simulated FC. In that case, in order to obtained simulations similar to empirical fMRI BOLD data, the dynamics of each node was binarized by putting all excitation to one and the remaining into 0 and then we convolved the binarized time-series with the standard hemodynamic response function to model the neurometabolic coupling [64]. As for empirical data, simulated FC was computed as the zero-lag Pearson correlation coefficient between the simulated BOLD timeseries of each pairs of brain regions. Furthermore, we explored topological properties of the original weighted structural network as well as the binarized structural network for which the correlation with the empirical FC was maximal. Specifically, we computed the network density, quantifying the proportion of edges that actually exist; the smallworld propensity, which is a robust measure of smallworldness (the presence of high local clustering versus short average path length) [68]; and modularity, which quantifies the level of segregation of a network into subgroups of densely connected nodes [12]. The smallworld propensity was computed using the code of the original publication (https://complexsystemsupenn.com/codedata) [68], and the others measures using the Brain Connectivity Toolbox (http://www.brain-connectivity-toolbox.net) [83].

### The perceptron

We used a simple perceptron model with one hidden layer in order to perform a multiclass classification task, here represented by handwritten digits from the MNIST benchmark dataset [27]. MNIST consists in a large set of 28×28 gray-scale images representing digits from 0 to 9 (60,000 training and 10,000 test examples). All images were normalized between 0 and 1. We trained networks with 3 layers: one input layer with 784=28×28 units, one unit for each pixel of the images; one hidden layer of 300 units, with a sharp sigmoid as activation function; and one output layer with one unit per digit (10 in total), with the softmax function as activation function.

The hidden and output layers are determined each by a connectivity matrix *W* and a bias vector *b*. An input image is first flattened to a vector, *X*, and propagated through the hidden layer *L_h_*(*X*) = *σ_h_*(*W_h_X*+*b_h_*), where *σ_h_* is the sigmoid function, *σ_h_*(*x*) = 1*/*(1 + *e*^−50^*^x^*). Here we chose a sharp sigmoid, i.e., with a scaling factor of 50, to mimic the threshold behavior of excitable networks. Then, the signal propagated to the output layer *L_o_* (*X*) = *σ_o_* (*W_o_ L_h_* (*X*) + *b_o_*), where *σ_o_* represents the softmax function, 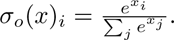

We optimized the networks using the crossentropy loss function which is minimized using stochastic gradient descent (SGD) over 10 epochs. All connection weights (*W*) are constrained to be non-negative. Furthermore, the weights to the hidden layer (*W_h_*) are regularized using *l*_1_ and *l*_2_ norms (*l*_1_=0.01, *l*_2_=0.0001). All simulations were performed using the machine learning library Tensorflow [1].

Similar to the SER model, we expect a network-binarizing capacity of the model at a threshold value of zero according to the sharp sigmoid activation function used. Moreover, given the nature of the input images, they approximatly mimic excitation patterns with binary activity, i.e., 0 (background) or 1 (drawing). Following the same reasoning as before, a pixel (or excitation) can propagate throught a given hidden unit, if the sum of the connection weight plus the bias exceed zero. Consequently, we can threshold and binarize the matrix *W_h_* column by column according to the bias vector *b_h_*, i.e.,

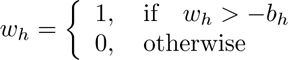

This thresholding and binarization also imply the removal of the bias vector.

## Data availability

Synthetic graphs were generated using the Matlab Tools for Network Analysis (original version 2006-2011, http://strategic.mit.edu/downloads.php?page=matlab_networks). The empirical brain network connectivity dataset is freely and publicly available on the Zenodo repository (https://zenodo.org/records/2872624) [43], while the human brain data on criticality was provided by Dr Enzo Tagliazucchi. The MNIST dataset is available on the Tensorflow library (https://www.tensorflow.org/datasets/catalog/mnist?hl=fr) [1].

## Code availability

Computational experiments were mostly performed in Matlab R2022a, while artificial neural networks were explored using Python 3.10.12. Topological graph properties were computed using the Brain Connectivity toolbox (version 2017-15-01, http://www.brain-connectivity-toolbox.net), except for the smallworld propensity which was computed using the original code (https://complexsystemsupenn.com/codedata). Additionally, we explored the computational advantages of the framework using three toolboxes: the Brain connectivity Toolbox (version 2017-15-01, http://www.brain-connectivity-toolbox.net), NetworkX (version 3.2.1, https://networkx.org) and igraph (version 0.11.3, https://igraph.org). Artificial neural networks were explored using the Tensorflow library (version 2.15.0, https://www.tensorflow.org/). All others simulations and analysis were done using custom matlab codes. Minimal code for exploring the framework developed in this study can be found on the Zenodo repository (provide link upon acceptance).

## Acknowledgements

This work was supported by funding from the Deutsche Forschungsgemeinschaft (DFG, German Research Foundation) - SFB 936 - 178316478 - A1 (C.C.H.) & Z3 (C.C.H. and A.M.), SPP2041 - 313856816 - HI1286/7-1 (C.C.H.) and TRR 169 - 261402652 - A2 (C.C.H.), and the European Union’s Horizon 2020 Framework Programme for Research and Innovation under Specific Grant Agreements 785907 (Human Brain Project SGA2 SGA3, C.C.H.). The funders had no role in study design, data collection and analysis, decision to publish, or preparation of the manuscript. We thank Dr Enzo Tagliazucchi for providing human brain data on criticality, Kayson Fakhar for helpful discussions on the figures design and Dr Wilhelm Braun for critical review of the manuscript.

## Ethics declarations

The authors declare no competing interests.

## Supplementary information

### FHN model threshold

The phase plane of the FHN model is characterized by the nullclines of the system, namely when *f* (*u, v*) = *g*(*u, v*) = 0, and are given by

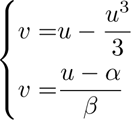

The intersection of the nullclines represents the fixed-points (*u*^∗^*, v*^∗^), they are given by the solutions of

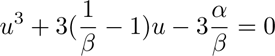

The solutions of this equation can be derived by the Cardano’s method

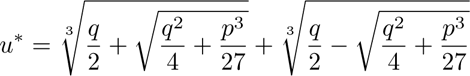

where 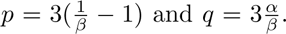 For the FHN to be in an excitable regime, *u*^∗^ must be lower than the local minimum of the N-shaped nullcline, namely *u^m^*, where *u^m^* is the solution of 1 − *u*^2^ = 0, i.e., *u^m^* = −1. We denoted *d* the distance of the fixed point to the unstable branch of the N-shaped nullcline *u^t^*, *d* = *u^t^* − *u*^∗^, where *u^t^* is one of the solution of

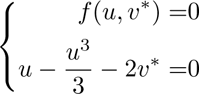

To compute the energy E transferred by a spike through the network, we resorted to numerical experiments, using a toy example composed of a pair of nodes connected with a connection of weight one (Fig S1). Varying the *α* parameter, we effectively vary the propagation threshold, by varying indirectly *d*. By initiating a spike in one of the two nodes and varying the coupling strength parameter *γ*, we can detect the critical *γ* (minimal value) that effectively propagates the spike to the second node, i.e., the model threshold. The model threshold is approximately a linear function of *d* (Fig S1b). However, we observed noticeable differences such as the presence of an offset as well as a slightly curved slope of the relationship between *d* and the model threshold. Such discrepancies can be explained by the highly non-linear nature of the FHN model, where the spike propagation is diffuse and filtered. Moreover, the offset can be intuitively understood. When *d* → 0 a critical point emerges where the system undergoes a Poincaŕe–Andronov–Hopf bifurcation, and therefore the system needs stronger than expected input force to escape this point, otherwise we observe damped small-amplitude oscillations returning to the fixed point. Plugging this quantity into the condition on network weights for spike propagation

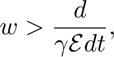

we can thus effectively modulate the possible set of links that can propagate the activity *via* theirs weights by varying the coupling parameter *γ* (Fig S1b).

**Figure S1.**
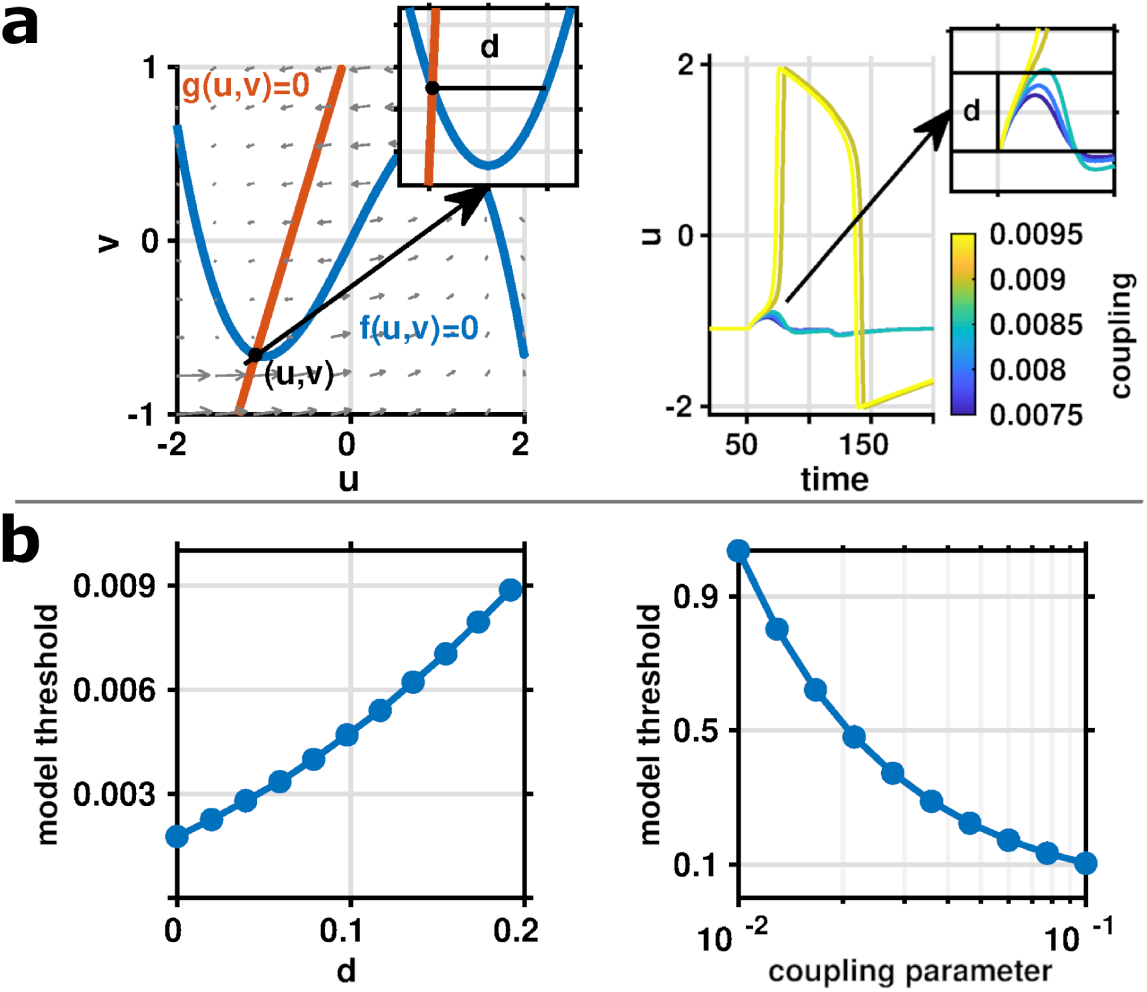
FHN model threshold. (a) Phase plane, where blue and red curves represent the nullclines (left), and propagation of a spike to a connected neighbor as a function of the coupling strength value (right). (b) The model threshold extracted from the exemplar setting as a function of *d* (left) and as a function of the coupling parameter *γ*, when *α* is fixed to 0.7 (right).

### Sytematic exploration

To investigate the role of topology and weight distribution in the formation of excitation patterns within networks, we considered three different types of undirected benchmark graphs: random, scale-free, and modular networks. The random graph was the classical Erdős-Ŕenyi (ER) model [29], the scale-free graph was the widely employed Barabási-Albert (BA) model [6], and the modular graph was a composition of a small number of random communities sparsely interconnected. We explored the robustness of the predictions across various network realization, size (*N* = 100 and 1000 for the SER model and 200 for the FHN model) and average degree (*k* = 4, 8 and 16). For random modular graphs, the number of modules is sampled uniformly between 2 and 8. All synthetic graphs were generated using the Matlab Tools for Network Analysis (http://strategic.mit.edu/downloads.php?page=matlab_networks) [13].

Subsequently, these binary networks are transformed into weighted networks by assigning weights to each connection following three typical distributions: a continuous uniform distribution between 0 and 1, a normal distribution with mean 0.5 and standard deviation 1/6, and a log-normal distribution with mean -1.5 and standard deviation 1. The parameters of the distributions are chosen such that the values lie approximately in the interval [0 1]. Negative values are discarded by taking the absolute value.

Predictions generally work well for sparse graphs and appear to be more influenced by the distribution of the weights than by the underlying network topology (Figs S2 and S3). See Figs S4 and S5 for the results on larger graphs. Predictions from the FHN model deviate for dense networks (Fig S3). However, the matching (correlation) between FC computed from weighted graphs and the ones from binary graphs remains high, implying that, although the predictions deviate, there still exists a correspondence. Predictions from the SER model work for a larger configuration range compared to the FHN model, only the predictions for dense networks ‘and’ large threshold values deviate from the theory (Fig S2).

**Figure S2.**
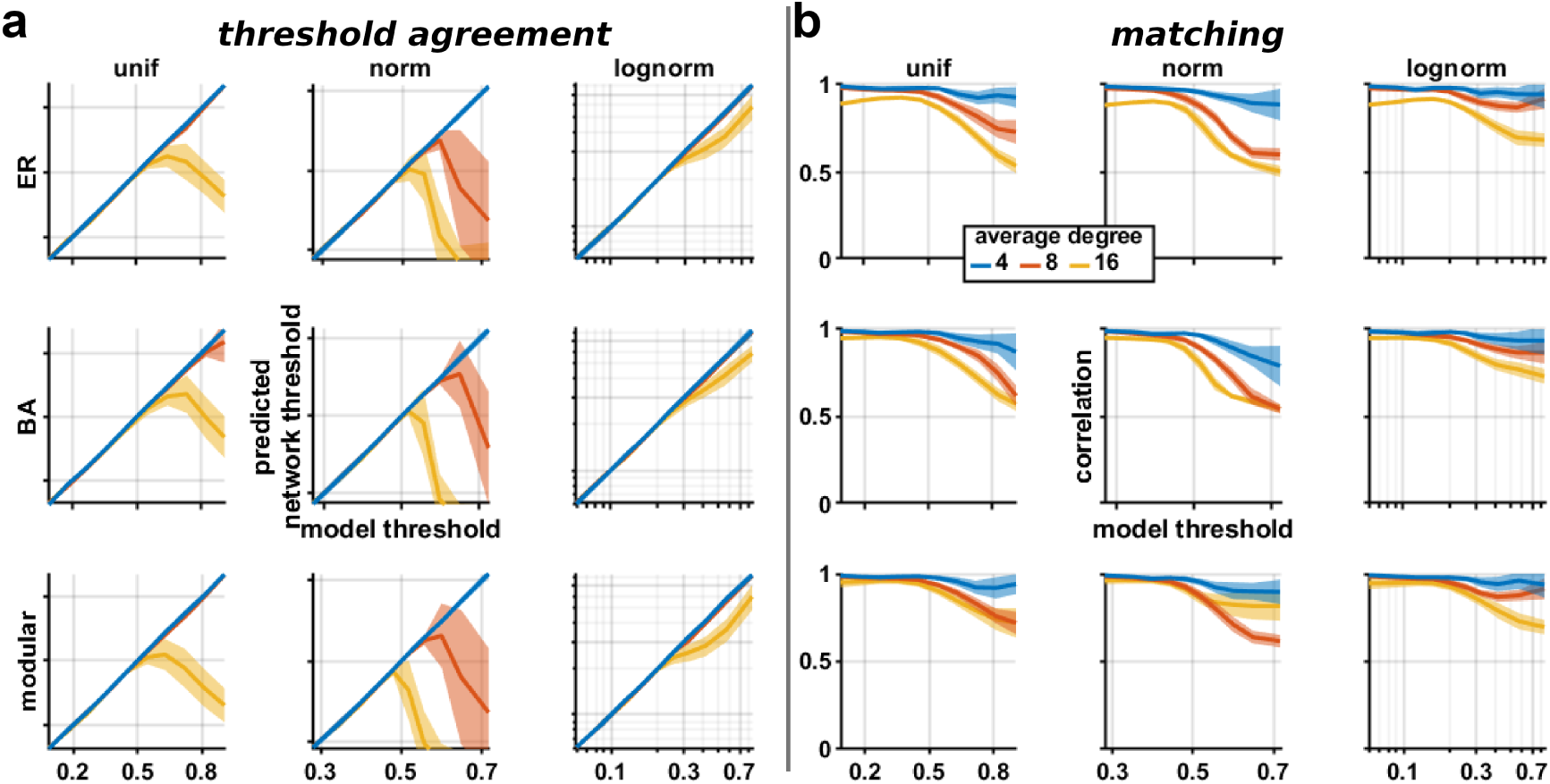
SER model exploration. (a) Threshold agreement, i.e., the predicted network threshold (y-axis) for which the corresponding FC is maximally correlated with the FC from the SER model running on the weighted version of the network, as a function of the model threshold (x-axis). (b) Matching, i.e., the corresponding maximal correlation between the FC patterns. Rows represent typical network topologies (random ER, scale-free BA, and modular), while columns correspond to different weight distributions. Curves and shaded areas represent the average and standard deviation over 50 network realizations and colors code for different average degree. Network size was fixed to 100 nodes.

### Number of excited neighbors

To better understand the discrepancies between theory and simulations, we first verify our main assumption which is that excitations are triggered mainly by a single excited neighbor. For that purpose, using the SER model and across the simulations on weighted networks, we computed the proportion of excitations originating from one or two excited neighbors. We observed that, when the model threshold increases, the proportion of nodes excited by one neighbor decreases while the proportion of nodes excited by two neighbors increases (Fig S6). Of note, these quantities appear largely insensitive to the network topology and network size and only sightly affected by the weight distribution. The increased proportion of excitations triggered by two neighbors appears to match the threshold discrepancies, as this quantity coincides with the departure of our prediction for the simulations of high threshold values. Of note, nearly all excitations are explained by excitation coming from at most two neighbors.

### Computational advantages

Thresholded and binarized networks present computational advantages both in terms of memory and execution time, here we show the time execution of basic matrix operations and topological caracteristics as well as the memory requirement of the networks and this through diverse software environments (Fig S7). To determine these computational aspects, we used random weighted networks with an uniform weight distribution and a fixed average degree of 20 and varying number of nodes (from 500 to 20,000). Subsequently, these networks were thresholded at different values (0.25, 0.5 and 0.75). We used 3 classic toolboxes: the Brain Connectivity Toolbox (http://www.brain-connectivity-toolbox.net) [83] developped for matlab (version 9.12.0 (R2022a), Natick Massachusetts: The MathWorks Inc. https://www.mathworks.com), NetworkX (https://networkx.org/) [44] and igraph (https://igraph.org/) [25] both developped in python (Python Software Foundation. Python Language Reference, version 3.11.5. http://www.python.org). All the experiments were performed on an AMD EPYC 75F3 server with 32 CPU cores at 2.95 GHz with 1 TB RAM. Simulations were repeated 5 times and the median execution times are shown.

**Figure S3.**
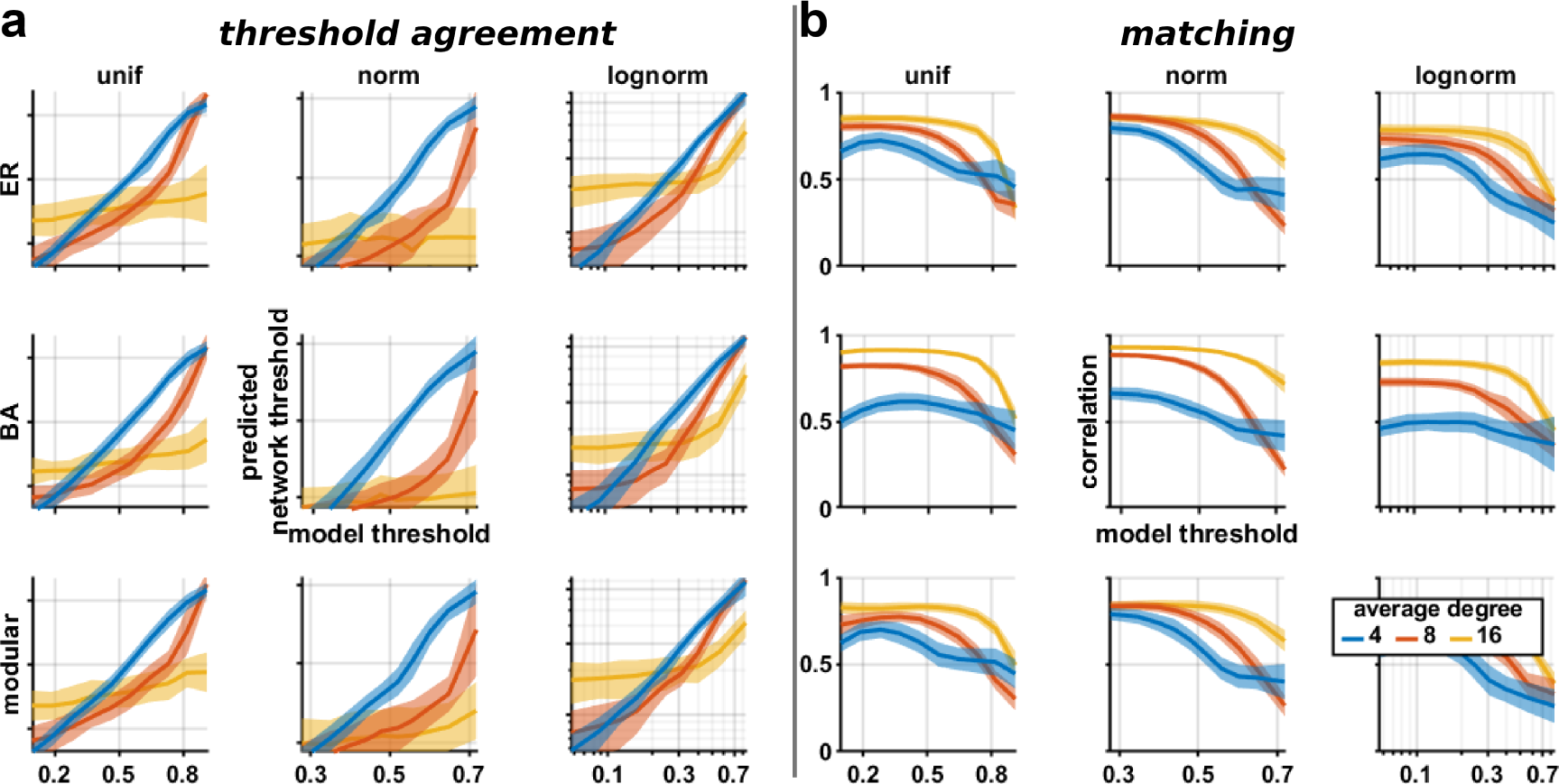
FHN model exploration. Same as Fig S2 but for the FHN model.

**Figure S4.**
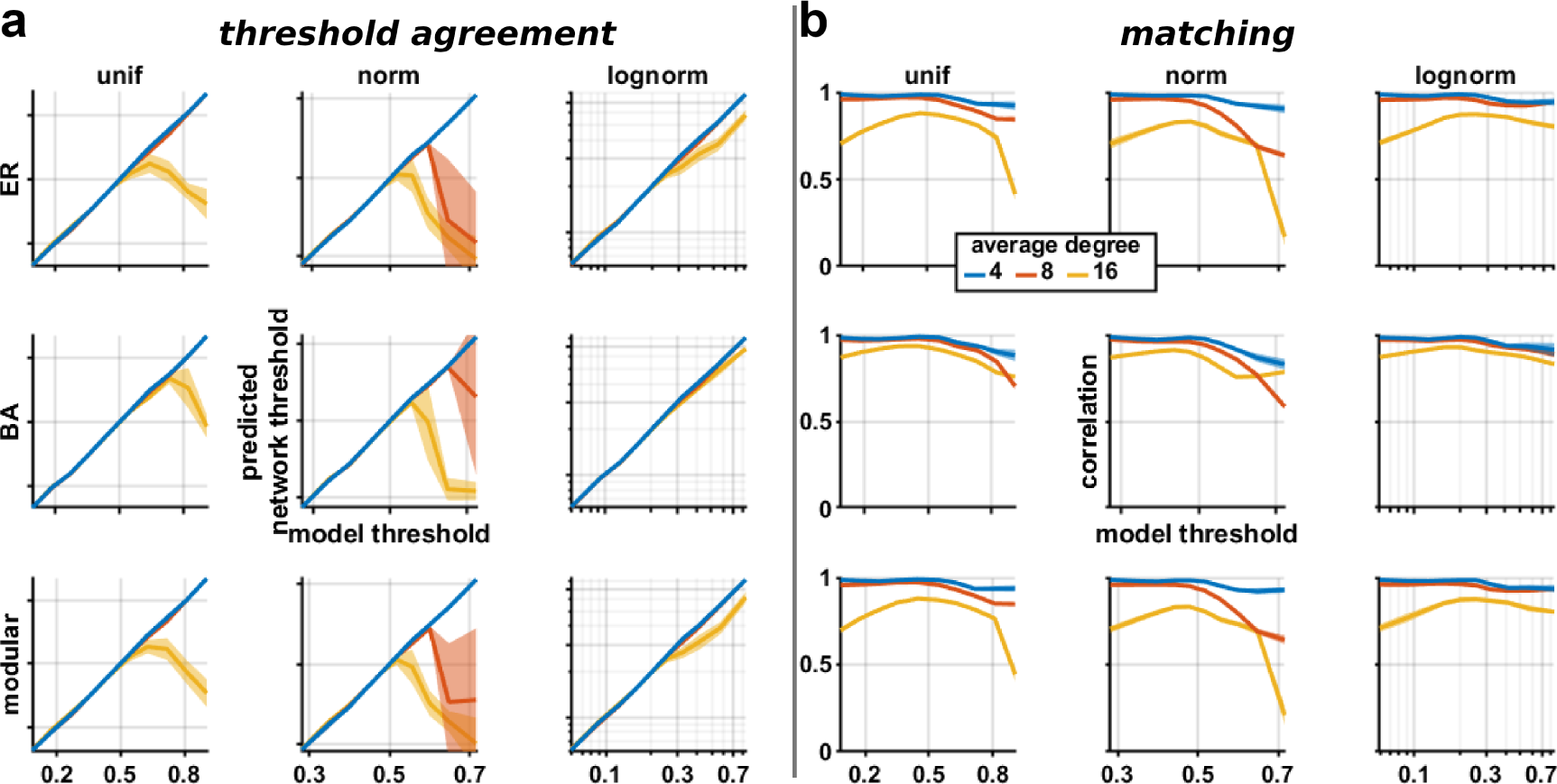
SER model exploration. Same as Fig S2 but for network size fixed to 1000 nodes.

**Figure S5.**
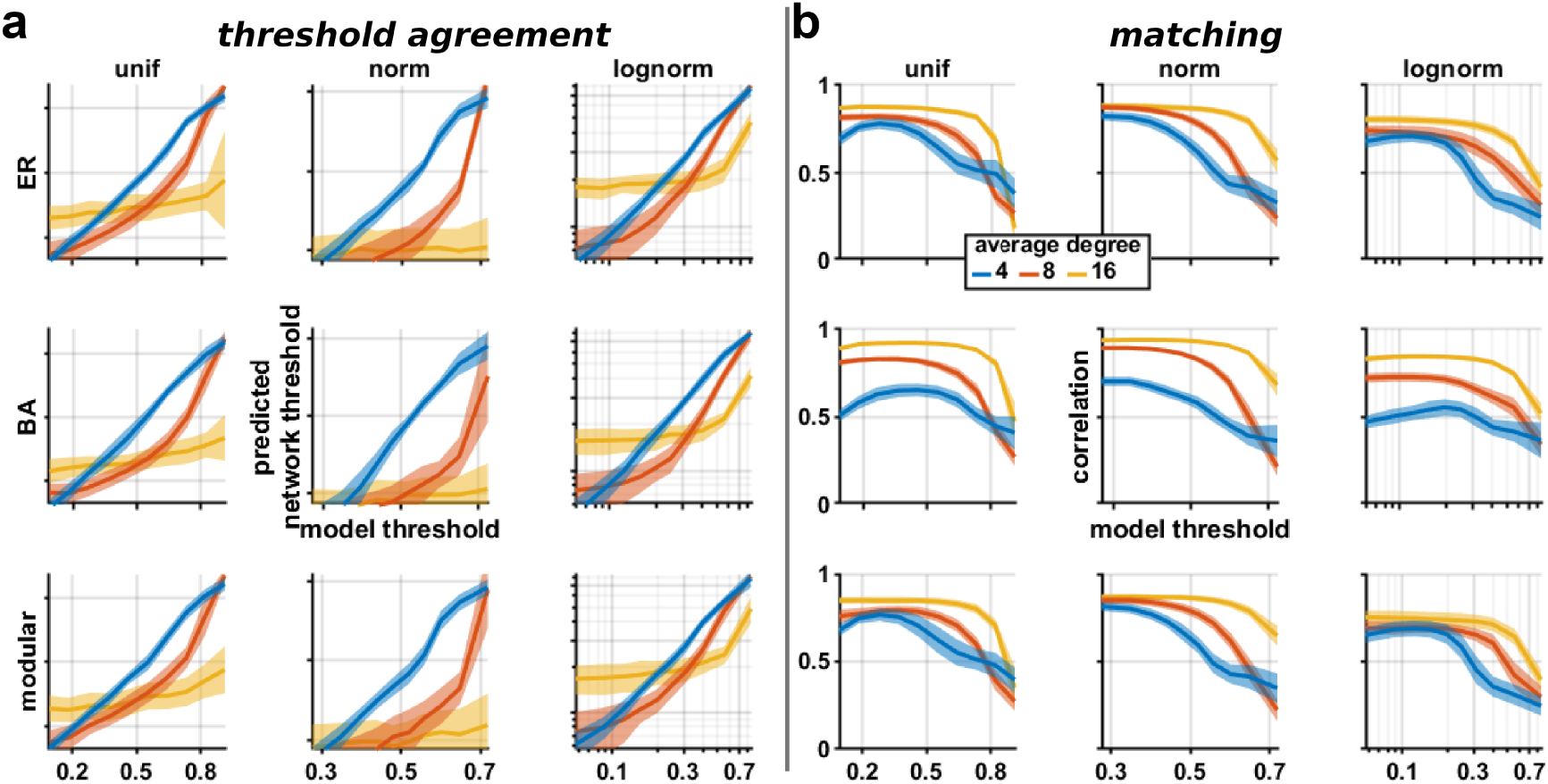
FHN model exploration. Same as Fig S3 but for network size fixed to 200 nodes.

As basic matrix operation we chose the matrix square operation, additionaly we computed topological properties including the shortest path from a single source node, the average clustering, the centrality based on eigenvector, and the modularity using greedy maximization. While we observed variations depending on the software and the measure used, overall, using binarized networks reduces the memory footprint as well as the time to execute mathematical operations (Fig S7).

### Brain criticality

We also explored key dynamical properties of resting-state fMRI activity. It has been shown that the brain at rest is near a critical point of a second order phase transition [21, 94]. Close to the critical point, the dynamics exhibit characteristics spatiotemporal patterns, especially scaling properties, such as divergence of the correlation length and the anomalous scaling of the correlation fluctuations.

Here, we first reproduced previous seminal findings of critical brain dynamics behavior captured by the conventional SER model [45]. Subsequently, we compared the results with those obtained from thresholded and binarized versions of the structural brain connectome (Figs S8 and S9). In particular, we computed two robust features exhibited by the spontaneous activity of human brain networks [33], the divergence of the correlation length and the quasi constant variance of the short-term pairwise correlations. All analyses were performed following the methods of the original paper and we refer the reader to it for further details [45].

### Generalization

While the main results are based on a simple setting, we show here that the framework is flexible and can therefore generalize to various alternative scenarios.

First, the threshold equivalence is also applicable for diverse network configurations, such as directed networks (Fig S10) and when the distribution of the weights depends on the underlying topology (Fig S11). Correlation between weights and network topology has been observed empirically [7]. Here, we generated networks where weights are proportional to the degree product. For that, we sampled weights (the number of samples corresponds to the number of connections) according to the given distribution. Then, we computed the degree product of all the connections (of the initially binary network). And finally, we mapped each weight according to the degree product, i.e., the lowest weight to the lowest degree product and so forth.

**Figure S6.**
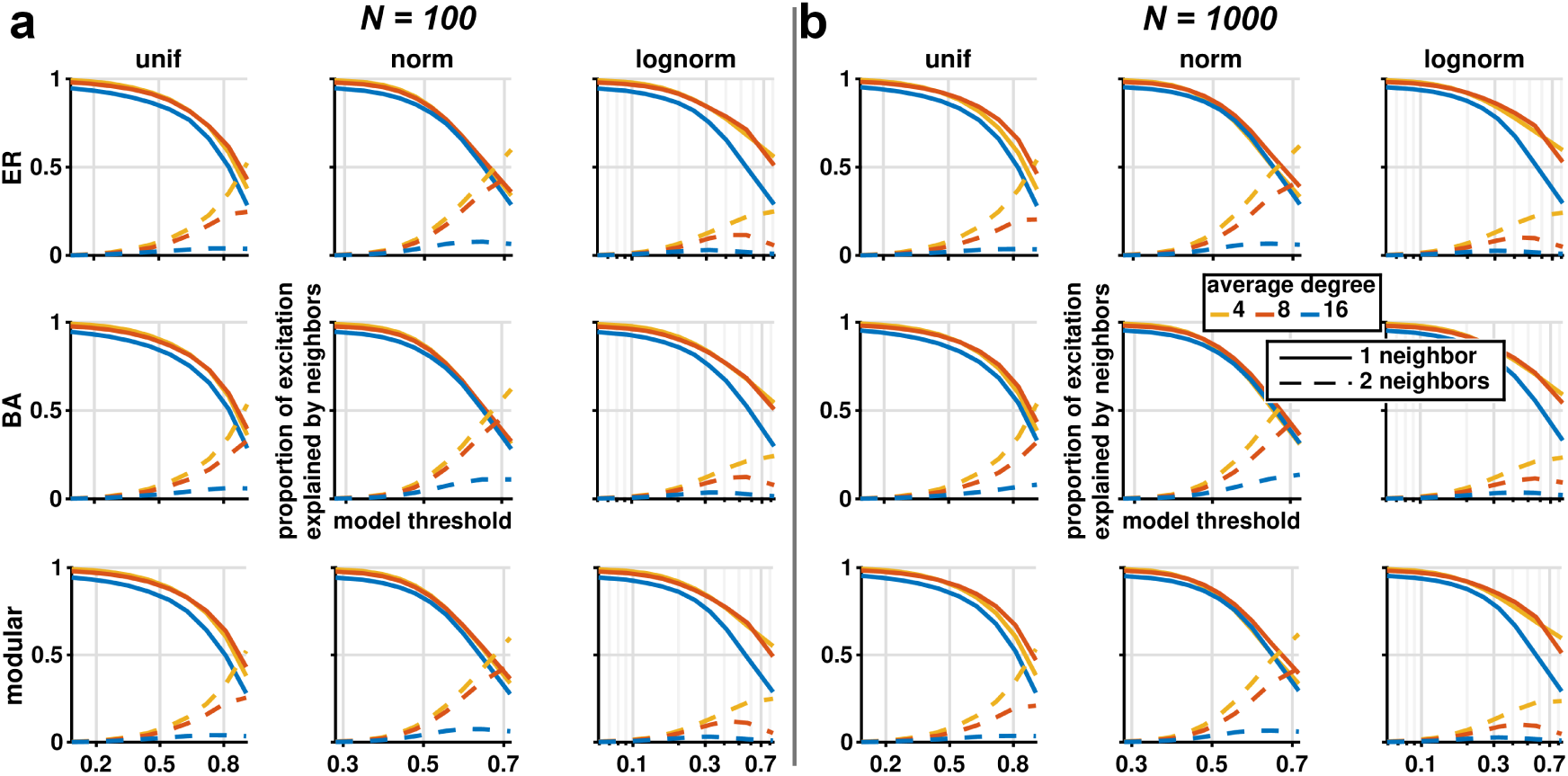
Proportion of excitation explained by neighbors. Proportion of excitations (y-axis) originating from one (solid curves) or two (dashed curves) excited neighbors as a function of the model threshold (x-axis) for the SER model. Rows represent typical network topologies (random ER, scale-free BA, and modular), while, columns correspond to different weight distributions. Curves represent the average over 50 network realizations and colors code for different average degree. Network size was fixed to 100 (a) and 1000 (b) nodes.

Second, the framework also generalizes to variations in the dynamical model, such as when network nodes have heterogeneous threshold values, e.g., when using a relative threshold (Fig S12). In contrast to the main SER formalism (absolute threshold), we used a relative threshold *T*, i.e., a node gets excited when Σ_N_*_E_ w > sT* where N*_E_* being the number of excited neighbors and *s* the strength of the node [50]. Consequently, each node has its own threshold value. When considering heterogeneous thresholds, the framework is virtually unchanged as long as a proper normalization is applied to the structural network. In the case of a relative threshold, it is equivalent to normalize each row of the structural network by its nodal strength (also called a right stochastic matrix), and then apply the original threshold framework. The framework also works when the activation function is probabilistic, as long as it encodes a steep threshold (Fig S13). The original SER formalism uses implicitely a Heaviside function, we can use a more smooth activation function such as the sigmoid function, a node gets excited with probability 1*/*(1 + *e^-a(^*Σ_NE *w-T*_*^),^* where *a* regulates the steepness and is set to 10.

**Figure S7.**
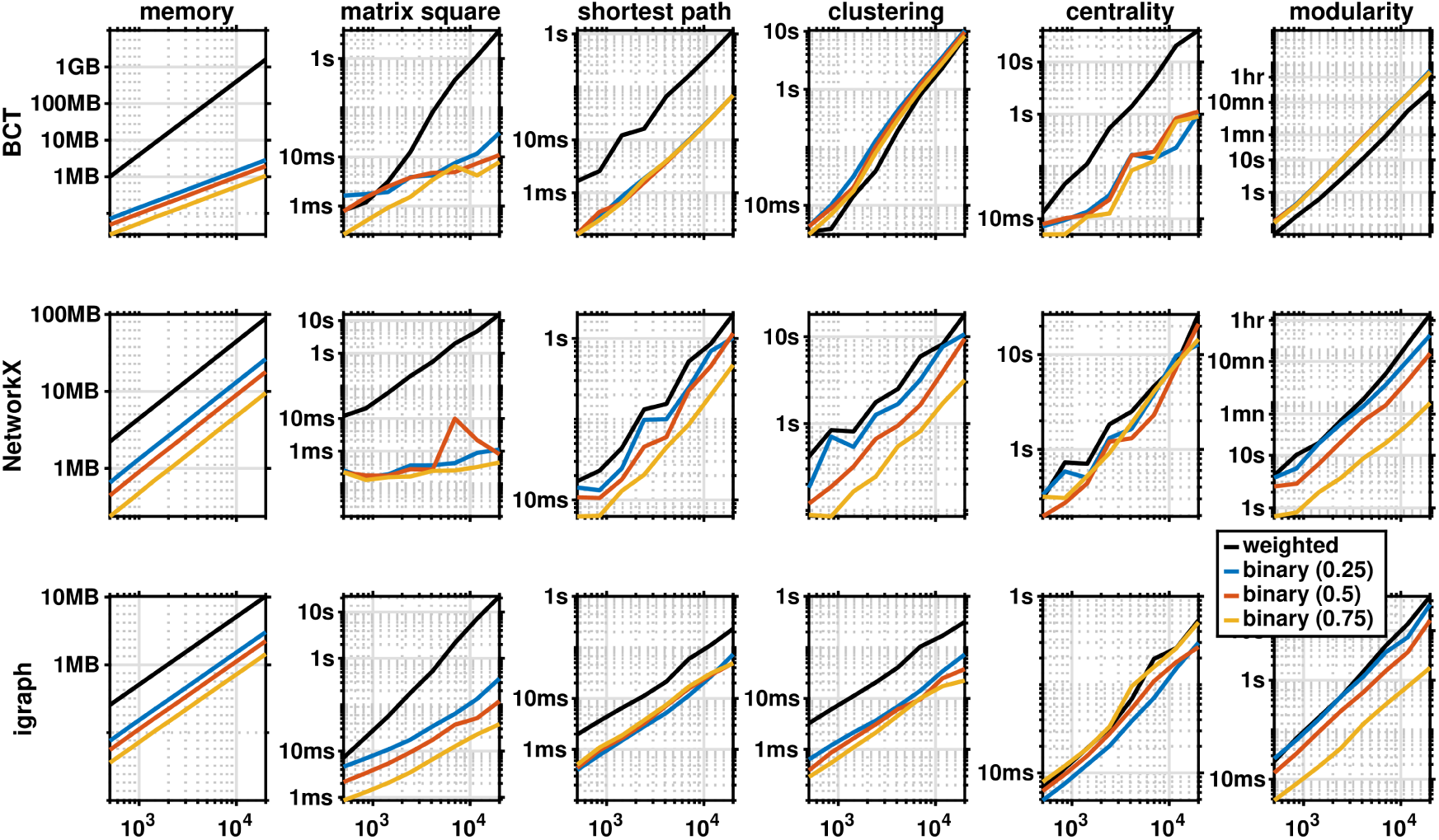
Computational advantages. Network memory footprint (in bytes) and execution time of basic matrix operation and topological properties as a function of the number of nodes (x-axis) and the network type (weighted or thresholded and binarized), and for different softwares (rows).

**Figure S8.**
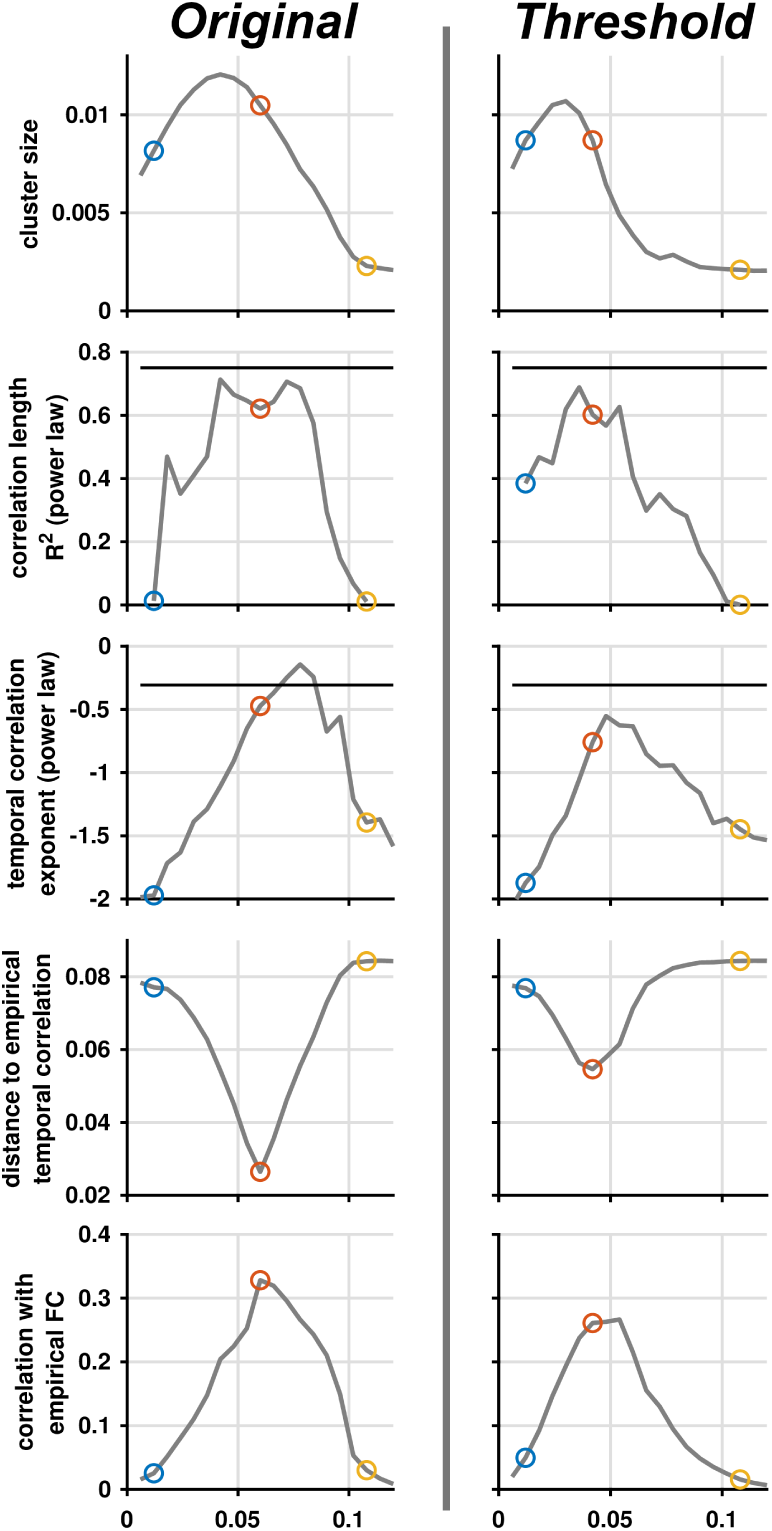
Exploration of brain criticality. Here we reproduce the original results from [45] about dynamical properties reminescent of criticality (first column) as well as the same quantities originating from our framework using thresholded and binarized networks (second column). The dynamical properties extracted are (from top to bottom): the average size of the second largest clusters of exicted nodes (Fig 1 in [45]); the explained variance (*R*^2^) of a power law fit of the correlation length (Fig S3 in [45]); the exponent of the power law fit of the temporal correlation fluctuations (Fig 3 in [45]); the distance of the temporal correlation fluctuations between the simulations and the empirical (Fig 3 in [45]); and finally the correlation between simulated and empirical FC (Fig S8 in [45]). All measures are as a function of the model threshold (first column) or the network threshold (second column). Three typical behaviors were highlighted by colored dots (below, around and above the critical regime), see Fig S9 for an illustration of the measures at these regimes.

**Figure S9.**
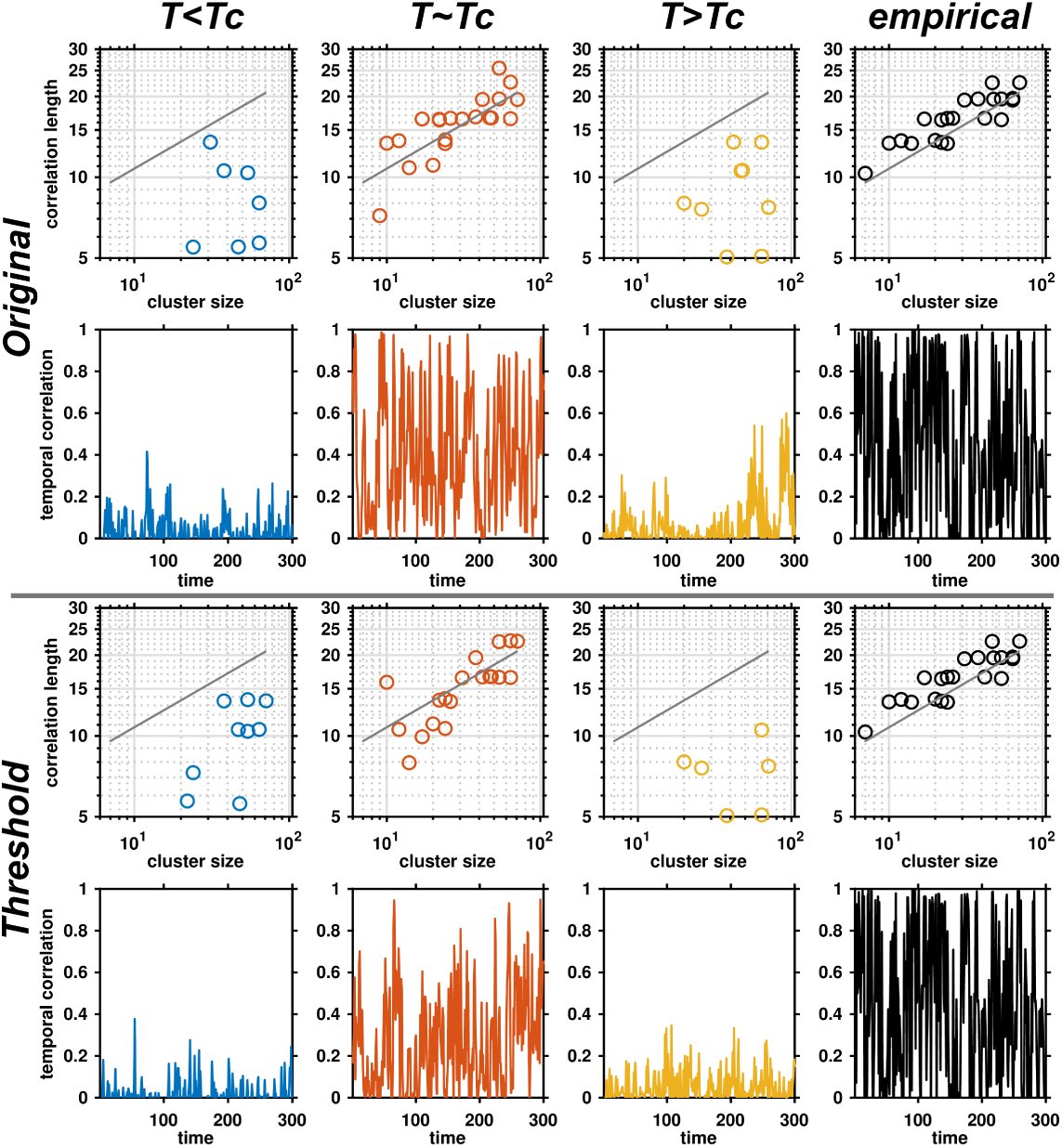
Correlation length and temporal correlation. (a) Examples of correlation length scaling as a function of the cluster size (Fig 2 in [45]) and of short-term correlation in the SER model at 3 dynamical regimes, below, around and above the critical point *T_c_* (Fig 3 in [45]), as well as for the human brain data (empirical). (b) Same as (a) but for the thresholded and binarized version of the SER model.

**Figure S10.**
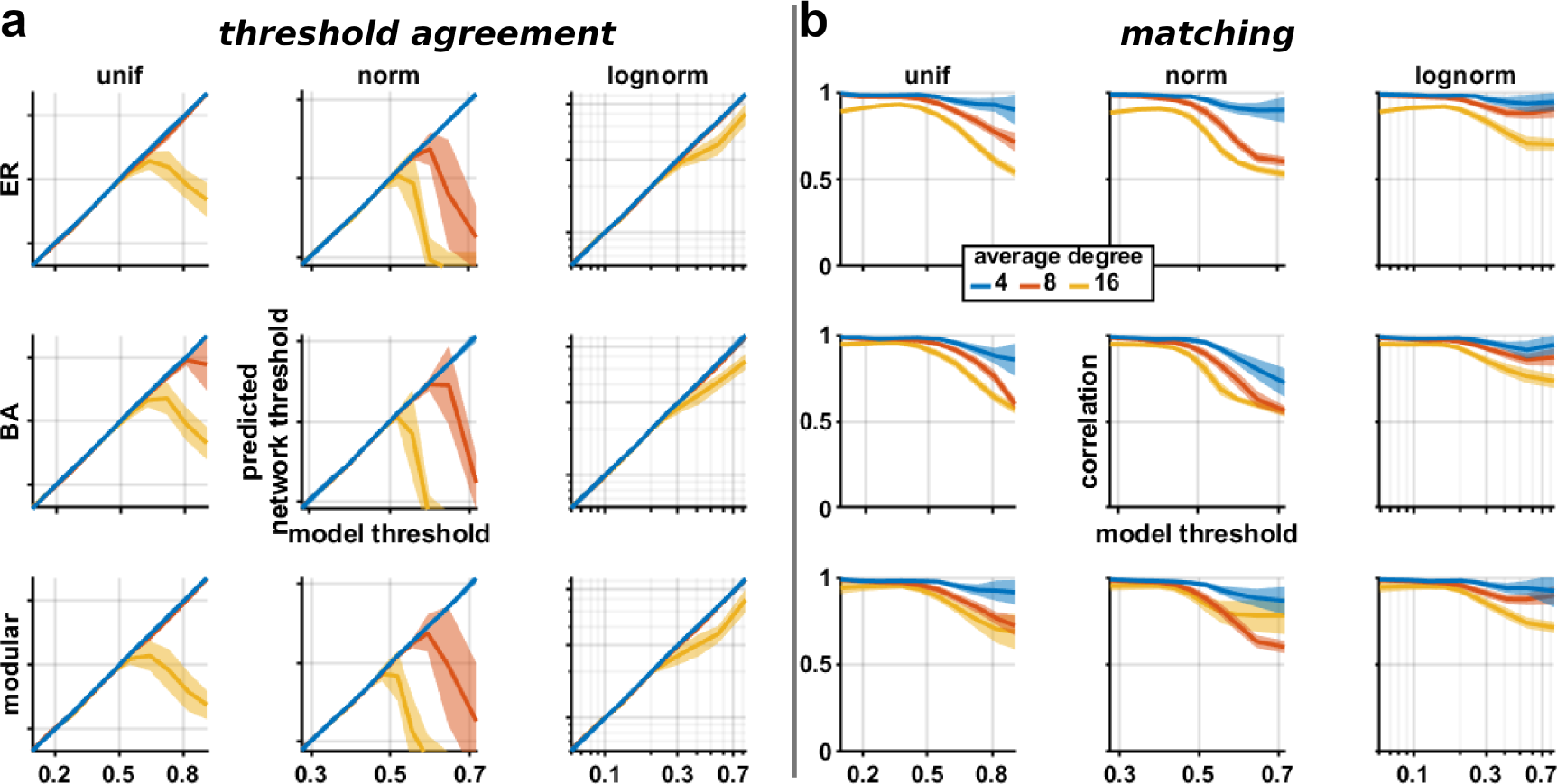
SER model exploration on directed networks. Same as Fig S2 but for directed network.

**Figure S11.**
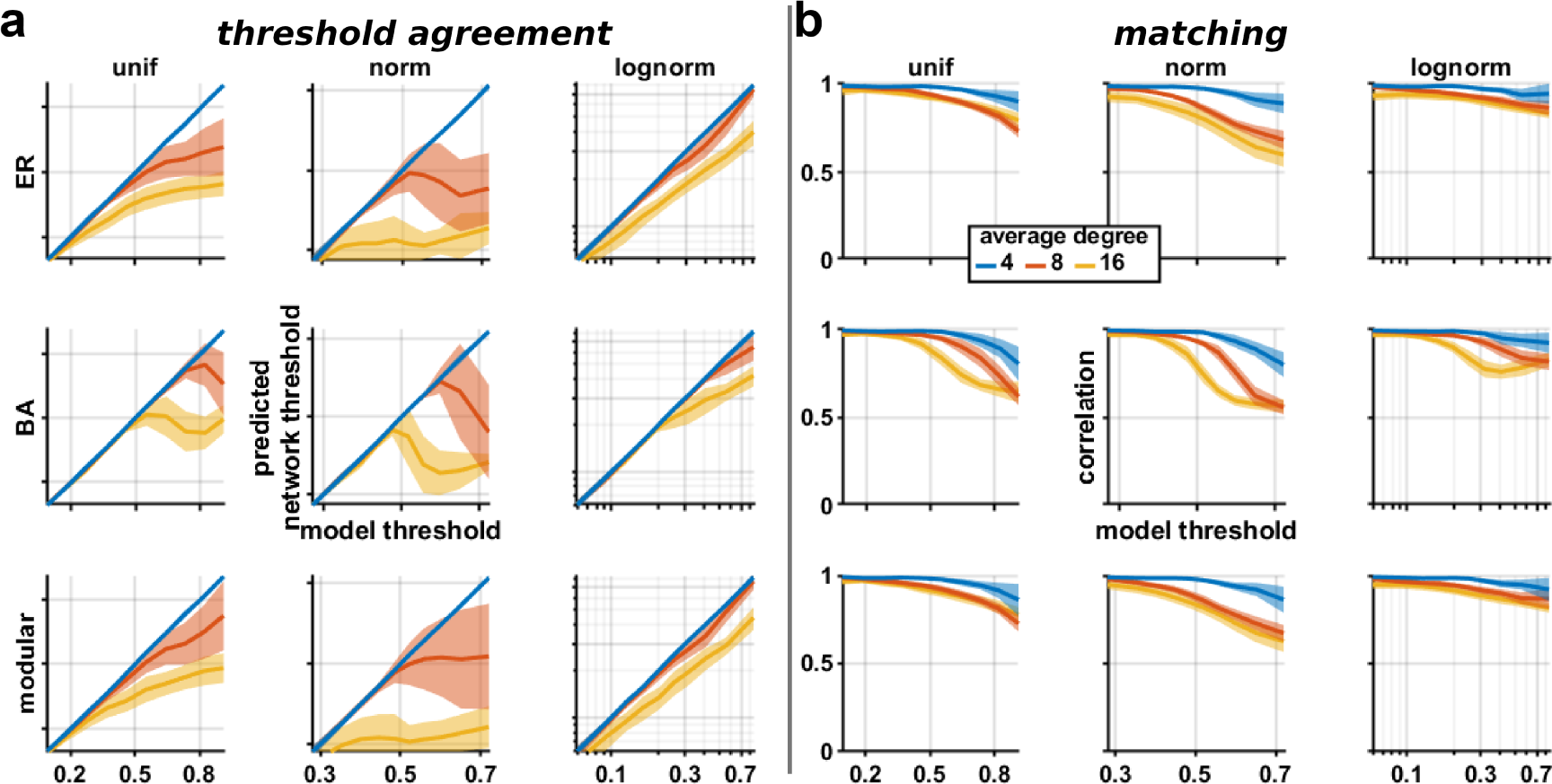
SER model exploration when network weights depend on network topology. Same as Fig S2 but for networks where weights are correlated to network degree.

**Figure S12.**
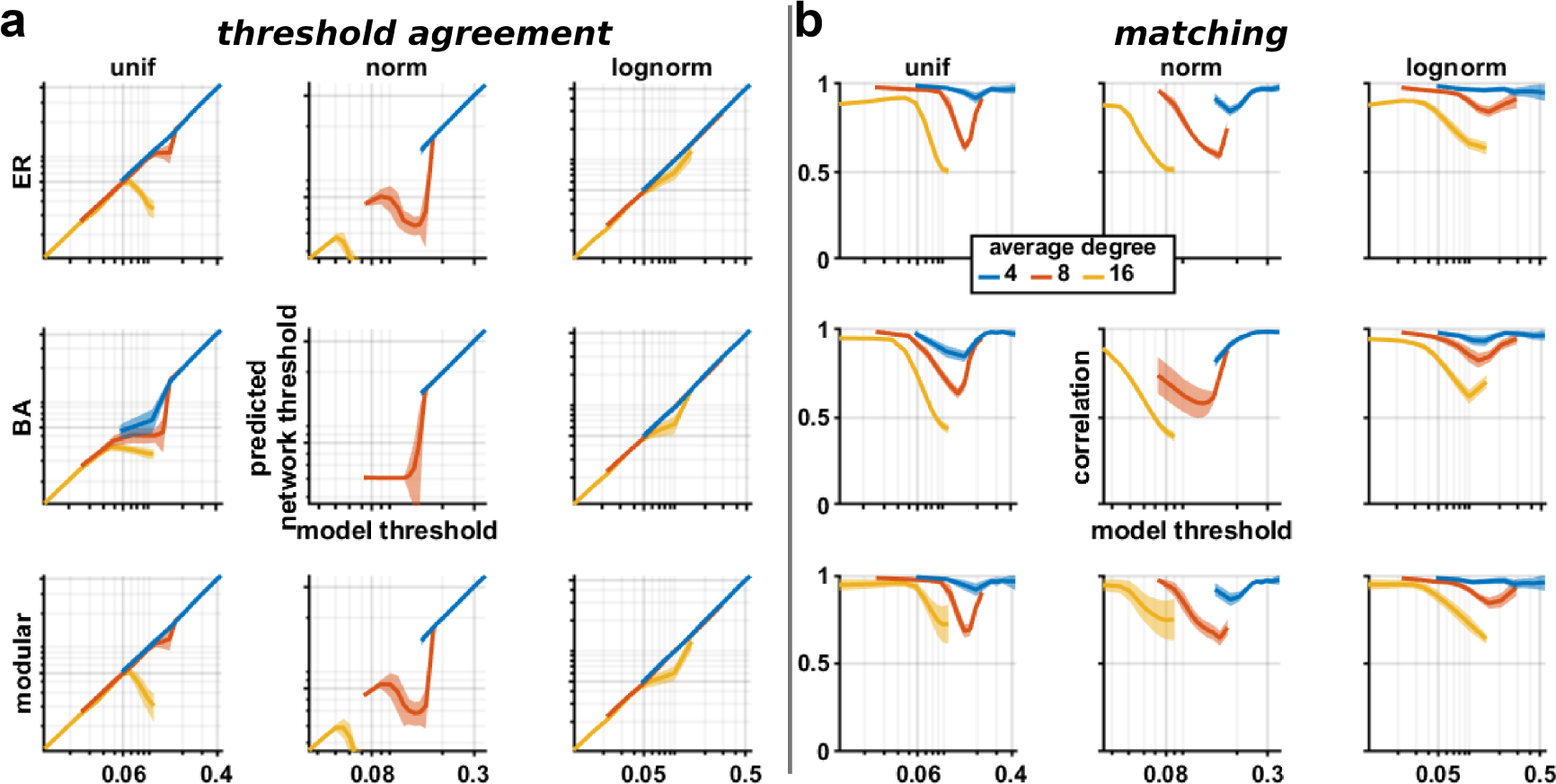
SER model exploration when using a relative threshold. Same as Fig S2 but using a relative threshold.

**Figure S13.**
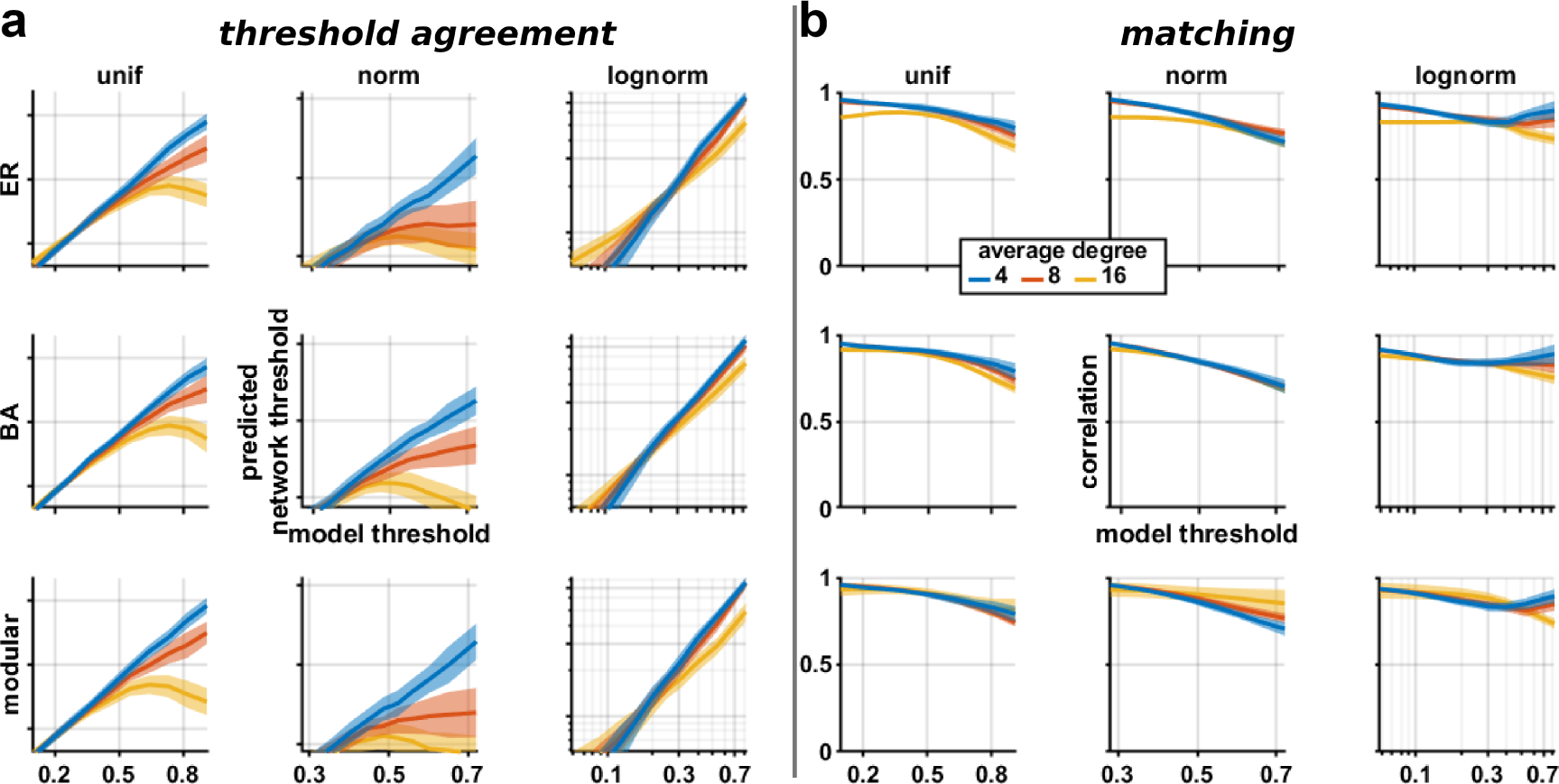
SER model exploration when using probabilistic activation function. Same as Fig S2 but using a sigmoid as probabilistic activation function.

## References

1. Martín Abadi, Ashish Agarwal, Paul Barham, Eugene Brevdo, Zhifeng Chen, Craig Citro, Greg S. Corrado, Andy Davis, Jeffrey Dean, Matthieu Devin, Sanjay Ghemawat, Ian Goodfellow, Andrew Harp, Geoffrey Irving, Michael Isard, Yangqing Jia, Rafal Jozefowicz, Lukasz Kaiser, Manjunath Kudlur, Josh Levenberg, Dandelion Mańe, Rajat Monga, Sherry Moore, Derek Murray, Chris Olah, Mike Schuster, Jonathon Shlens, Benoit Steiner, Ilya Sutskever, Kunal Talwar, Paul Tucker, Vincent Vanhoucke, Vijay Vasudevan, Fernanda Viégas, Oriol Vinyals, Pete Warden, Martin Wattenberg, Martin Wicke, Yuan Yu, and Xiaoqiang Zheng, TensorFlow: Large-scale machine learning on heterogeneous systems, 2015, Software available from tensorflow.org.

2. Ioannis E Antoniou and Eleni T Tsompa, Statistical analysis of weighted networks, Discrete dynamics in Nature and Society 2008 (2008), 1–16.

3. Alex Arenas, Albert Díaz-Guilera, and Conrad J Pérez-Vicente, Synchronization reveals topological scales in complex networks, Physical Review Letters 96 (2006), 114102.

4. Andrea Avena-Koenigsberger, Bratislav Misic, and Olaf Sporns, Communication dynamics in complex brain networks, Nature Reviews Neuroscience 19 (2018), 17–33.

5. Per Bak, Kan Chen, and Chao Tang, A forest-fire model and some thoughts on turbulence, Physics Letters A 147 (1990), 297–300.

6. Albert-László Barabási and Reka Albert, Emergence of scaling in random networks, Science 286 (1999), 15551–15555.

7. Alain Barrat, Marc Barthélemy, Romualdo Pastor-Satorras, and Alessandro Vespignani, The architecture of complex weighted networks, Proceedings of the National Academy of Sciences of the U.S.A. 101 (2004), 3747–3752.

8. Marc Barthelemy, Alain Barrat, Romualdo Pastor-Satorras, and Alessandro Vespignani, Characterization and modeling of weighted networks, Physica A: Statistical Mechanics and its Applications 346 (2005), 34–43.

9. Danielle S Bassett and Olaf Sporns, Network neuroscience, Nature Neuroscience 20 (2017), 353–364.

10. Richard F Betzel and Danielle S Bassett, Specificity and robustness of long-distance connections in weighted, interareal connectomes, Proceedings of the National Academy of Sciences of the U.S.A. 115 (2018), E4880–E4889.

11. Sayak Bhattacharya and Pablo A Iglesias, The threshold of an excitable system serves as a control mechanism for noise filtering during chemotaxis, PLoS One 13 (2018), 1–16.

12. Vincent D Blondel, Jean-Loup Guillaume, Renaud Lambiotte, and Etienne Lefebvre, Fast unfolding of communities in large networks, Journal of Statistical Mechanics: Theory and Experiment 2008 (2008), P10008.

13. Gergana Bounova and Olivier de Weck, Overview of metrics and their correlation patterns for multiple-metric topology analysis on heterogeneous graph ensembles, Physical Review E 85 (2012), 016117.

14. Colin R. Buchanan, Mark E. Bastin, Stuart J. Ritchie, David C. Liewald, James W. Madole, Elliot M. Tucker-Drob, Ian J. Deary, and Simon R. Cox, The effect of network thresholding and weighting on structural brain networks in the uk biobank, NeuroImage 211 (2020), 116443.

15. Edward Bullmore and Olaf Sporns, Complex brain networks: graph theoretical analysis of structural and functional systems, Nature Reviews Neuroscience 10 (2009), 186–198.

16. György Buzaáki and Kenji Mizuseki, The log-dynamic brain: how skewed distributions affect network operations, Nature Reviews Neuroscience 15 (2014), 264–278.

17. Joana Cabral, Morten L Kringelbach, and Gustavo Deco, Exploring the network dynamics underlying brain activity at rest, Progress in Neurobiology 114 (2014), 102–131.

18. Leila Cammoun, Xavier Gigandet, Djalel Meskaldji, Jean Philippe Thiran, Olaf Sporns, Kim Q Do, Philippe Maeder, Reto Meuli, and Patric Hagmann, Mapping the human connectome at multiple scales with diffusion spectrum mri, Journal of Neuroscience Methods 203 (2012), 386–397.

19. George T Cantwell, Yanchen Liu, Benjamin F Maier, Alice C Schwarze, Carlos A Serván, Jordan Snyder, and Guillaume St-Onge, Thresholding normally distributed data creates complex networks, Physical Review E 101 (2020), no. 6, 062302.

20. Yuhan Chen, Shengjun Wang, Claus C Hilgetag, and Changsong Zhou, Features of spatial and functional segregation and integration of the primate connectome revealed by trade-off between wiring cost and efficiency, PLoS Computational Biology 13 (2017), 1–37.

21. Dante R Chialvo, Emergent complex neural systems, Nature Physics 6 (2010), 744–750.

22. Oren Civier, Robert Elton Smith, Chun-Hung Yeh, Alan Connelly, and Fernando Calamante, Is removal of weak connections necessary for graph-theoretical analysis of dense weighted structural connectomes from diffusion mri?, NeuroImage 194 (2019), 68–81.

23. Gilberto Corso, Claudia Patricia Torres Cruz, Míriam Plaza Pinto, Adriana Monteiro de Almeida, and Thomas M Lewinsohn, Binary versus weighted interaction networks, Ecological Complexity 23 (2015), 68–72.

24. Lee Cossell, Maria Florencia Lacaruso, Dylan R Muir, Rachael Houlton, Elie N Sader, Ho Ko, Sonja B Hofer, and Thomas D Mrsic-Flogel, Functional organization of excitatory synaptic strength in primary visual cortex, Nature 518 (2015), 399–403.

25. Gabor Csardi and Tamas Nepusz, *The igraph software package for complex network research*, InterJournal, Complex Systems 1695 (2006), 1–9.

26. Alessandro Daducci, Stephan Gerhard, Alessandra Griffa, Alia Lemkaddem, Leila Cammoun, Xavier Gigandet, Reto Meuli, Patric Hagmann, and Jean-Philippe Thiran, The connectome mapper: An open-source processing pipeline to map connectomes with mri, PLoS One 7 (2012), 1–9.

27. Li Deng, The mnist database of handwritten digit images for machine learning research [best of the web], IEEE Signal Processing Magazine 29 (2012), 141–142.

28. Rahul S Desikan, Florent Ségonne, Bruce Fischl, Brian T Quinn, Bradford C Dickerson, Deborah Blacker, Randy L Buckner, Anders M Dale, R Paul Maguire, Bradley T Hyman, Marilyn S Albert, and Ronald J Killiany, An automated labeling system for subdividing the human cerebral cortex on mri scans into gyral based regions of interest, NeuroImage 31 (2006), 968–980.

29. Paul Erdős and Alfréd Rényi, On the evolution of random graphs, Publications of the Mathematical Institute of the Hungarian Academy of Sciences 5 (1960), 17–61.

30. G Bard Ermentrout and Leah Edelstein-Keshet, Cellular automata approaches to biological modeling, Journal of Theoretical Biology 160 (1993), no. 1, 97–133.

31. Richard FitzHugh, Mathematical models of threshold phenomena in the nerve membrane, Bulletin of Mathematical Biology 17 (1955), 257–278.

32. Richard FitzHugh, Impulses and physiological states in theoretical models of nerve membrane, Biophysical Journal 1 (1961), 445–466.

33. Daniel Fraiman and Dante R. Chialvo, What kind of noise is brain noise: anomalous scaling behavior of the resting brain activity fluctuations, Frontiers in Physiology 3 (2012).

34. Karl J Friston, Functional and effective connectivity: a review, Brain connectivity 1 (2011), 13–36.

35. Lucas S Furtado and Mauro Copelli, Response of electrically coupled spiking neurons: a cellular automaton approach, Physical Review. E, Statistical, Nonlinear, and Soft Matter Physics 73 (2006), 011907+.

36. Guadalupe C Garcia, Annick Lesne, Marc-Thorsten Hütt, and Claus C Hilgetag, Building blocks of self-sustained activity in a simple deterministic model of excitable neural networks, Frontiers in Computational Neuroscience 6 (2012), 50.

37. Lea Goetz, Arnd Roth, and Michael Häusser, Active dendrites enable strong but sparse inputs to determine orientation selectivity, Proceedings of the National Academy of Sciences of the U.S.A. 118 (2021), e2017339118.

38. Carlos Henrique Gomes Ferreira, Fabricio Murai, Ana P C Silva, Martino Trevisan, Luca Vassio, Idilio Drago, Marco Mellia, and Jussara M Almeida, On network backbone extraction for modeling online collective behavior, 17 (2022), 1–36.

39. Joaquín Goñi, Martijn P van den Heuvel, Andrea Avena-Koenigsberger, Nieves Velez de Mendizabal, Richard F Betzel, Alessandra Griffa, Patric Hagmann, Bernat Corominas-Murtra, Jean-Philippe Thiran, and Olaf Sporns, Resting-brain functional connectivity predicted by analytic measures of network communication, Proceedings of the National Academy of Sciences of the U.S.A. 111 (2014), 833–838.

40. Mark Granovetter, The strength of weak ties: A network theory revisited, Sociological theory 1 (1983), 201–233.

41. Nicholas C Grassly and Christophe Fraser, Mathematical models of infectious disease transmission, Nature Reviews Microbiology 6 (2008), 477–487.

42. James M Greenberg and Stuart P Hastings, Spatial patterns for discrete models of diffusion in excitable media, SIAM Journal on Applied Mathematics 34 (1978), 515–523.

43. Alessandra Griffa, Yasser Alemán-Gómez, and Patric Hagmann, Structural and functional connectome from 70 young healthy adults, Dataset on Zenodo, 2019.

44. Aric A Hagberg, Daniel A Schult, and Pieter J Swart, Exploring network structure, dynamics, and function using networkx, Proceedings of the 7th Python in Science Conference (Pasadena, CA USA) (Gäel Varoquaux, Travis Vaught, and Jarrod Millman, eds.), 2008, pp. 11–15.

45. Ariel Haimovici, Enzo Tagliazucchi, Pablo Balenzuela, and Dante R Chialvo, Brain organization into resting state networks emerges at criticality on a model of the human connectome, Physical Review Letters 110 (2013), 178101.

46. Mike Hemberger, Mark Shein-Idelson, Lorenz Pammer, and Gilles Laurent, Reliable sequential activation of neural assemblies by single pyramidal cells in a three-layered cortex, Neuron 104 (2019), 353–369.e5.

47. Alan L Hodgkin and Andrew F Huxley, A quantitative description of membrane current and its application to conduction and excitation in nerve, The Journal of Physiology 117 (1952), 500–544.

48. Itay Hubara, Matthieu Courbariaux, Daniel Soudry, Ran El-Yaniv, and Yoshua Bengio, Binarized neural networks, Advances in neural information processing systems 29 (2016).

49. Marc-Thorsten Hütt, Claus C Hilgetag, and Marcus Kaiser, Network-guided pattern formation of neural dynamics, Philosophical Transactions of the Royal Society of London. Series B, Biological Sciences 369 (2014), 20130522.

50. Marc-Thorsten Hütt, Mitul Jain, Claus C Hilgetag, and Annick Lesne, Stochastic resonance in discrete excitable dynamics on graphs, Chaos, Solitons & Fractals 45 (2012), 611–618.

51. Ramakrishnan Iyer, Vilas Menon, Michael Buice, Christof Koch, and Stefan Mihalas, The influence of synaptic weight distribution on neuronal population dynamics, PLoS Computational Biology 9 (2013), 1–16.

52. Eugene M Izhikevich, Dynamical systems in neuroscience, Cambridge (Massachusetts): MIT Press, 2007.

53. Ho Ko, Sonja B Hofer, Bruno Pichler, Katherine A Buchanan, P Jesper Sjöström, and Thomas D Mrsic-Flogel, Functional specificity of local synaptic connections in neocortical networks, Nature 473 (2011), 87–91.

54. Stephan Krohn, Nina von Schwanenflug, Leonhard Waschke, Amy Romanello, Martin Gell, Douglas D. Garrett, and Carsten Finke, A spatiotemporal complexity architecture of human brain activity, Science Advances 9 (2023), eabq3851.

55. Łukasz Kuśmierz, Shun Ogawa, and Taro Toyoizumi, Edge of chaos and avalanches in neural networks with heavy-tailed synaptic weight distribution, Physical Review Letters 125 (2020), 028101.

56. Wei-Chung Allen Lee, Vincent Bonin, Michael Reed, Brett J Graham, Greg Hood, Katie Glattfelder, and R Clay Reid, Anatomy and function of an excitatory network in the visual cortex, Nature 532 (2016), 370–374.

57. Xiao Liu, David A. Leopold, and Yifan Yang, Single-neuron firing cascades underlie global spontaneous brain events, Proceedings of the National Academy of Sciences of the U.S.A. 118 (2021), e2105395118.

58. Yang-Yu Liu, Jean-Jacques Slotine, and Albert-László Barabási, Controllability of complex networks, Nature 473 (2011), 167–173.

59. Nikola T Markov, Mária M Ercsey-Ravasz, Ana Rita Ribeiro Gomes, Camille Lamy, Löıc Magrou, Julien Vezoli, Pierre Misery, Arnaud Falchier, René Quilodran, Marie-Alice Gariel, Jérôme Sallet, Răzvan Gămănut, Cyril Huissoud, Simon Clavagnier, Pascale Giroud, Dominique Sappey-Marinier, Pascal Barone, Colette Dehay, Zoltan Toroczkai, Kenneth Knoblauch, David C Van Essen, and Henry Kennedy, A weighted and directed interareal connectivity matrix for macaque cerebral cortex, Cerebral Cortex 24 (2014), 17–36.

60. Joachim Mathiesen, Luiza Angheluta, Peter T. H. Ahlgren, and Mogens H. Jensen, Excitable human dynamics driven by extrinsic events in massive communities, Proceedings of the National Academy of Sciences of the U.S.A. 110 (2013), 17259–17262.

61. Arnaud Messé, Habib Benali, and Guillaume Marrelec, Relating structural and functional connectivity in MRI: a simple model for a complex brain, IEEE Transactions on Medical Imaging 34 (2015), 27–37.

62. Arnaud Messé, Marc-Thorsten Hütt, and Claus C Hilgetag, Toward a theory of coactivation patterns in excitable neural networks, PLoS Computational Biology 14 (2018), 1–19.

63. Arnaud Messé, Marc-Thorsten Hütt, Peter König, and Claus C Hilgetag, A closer look at the apparent correlation of structural and functional connectivity in excitable neural networks, Scientific Reports 5 (2015), 7870.

64. Arnaud Messé, David Rudrauf, Habib Benali, and Guillaume Marrelec, Relating structure and function in the human brain: relative contributions of anatomy, stationary dynamics, and non-stationarities, PLoS Computational Biology 10 (2014), e1003530.

65. Arnaud Messé, David Rudrauf, Alain Giron, and Guillaume Marrelec, Predicting functional connectivity from structural connectivity via computational models using MRI: an extensive comparison study, NeuroImage 111 (2015), 65–75.

66. Yuchuan Miao, Sayak Bhattacharya, Marc Edwards, Huaqing Cai, Takanari Inoue, Pablo A Iglesias, and Peter N Devreotes, Altering the threshold of an excitable signal transduction network changes cell migratory modes, Nature Cell Biology 19 (2017), 329–340.

67. A.J. Morales, J. Borondo, J.C. Losada, and R.M. Benito, Efficiency of human activity on information spreading on twitter, Social Networks 39 (2014), 1–11.

68. Sarah Feldt Muldoon, Eric W Bridgeford, and Danielle S Bassett, Small-world propensity and weighted brain networks, Scientific Reports 6 (2016), 22057.

69. Mark Müller-Linow, Carsten Marr, and Marc-Thorsten Hütt, Topology regulates the distribution pattern of excitations in excitable dynamics on graphs, Physical Review E 74 (2006), 1–7.

70. Jinichi Nagumo, Suguru Arimoto, and Shuji Yoshizawa, An active pulse transmission line simulating nerve axon, Proceedings of the IRE 50 (1962), 2061–2070.

71. Karl-Heinz Nenning, Ting Xu, Alexandre R. Franco, Khena M. Swallow, Arielle Tambini, Daniel S. Margulies, Jonathan Smallwood, Stanley J. Colcombe, and Michael P. Milham, Omnipresence of the sensorimotor-association axis topography in the human connectome, NeuroImage 272 (2023), 120059.

72. Mark EJ Newman, Analysis of weighted networks, Physical Review E 70 (2004), 056131.

73. Jun nosuke Teramae, Yasuhiro Tsubo, and Tomoki Fukai, Optimal spike-based communication in excitable networks with strong-sparse and weak-dense links, Scientific Reports 2 (2012), 485.

74. Gabriel Koch Ocker, Krešimir Josić, Eric Shea-Brown, and Michael A Buice, Linking structure and activity in nonlinear spiking networks, PLoS Computational Biology 13 (2017), 1–47.

75. Seung W Oh, Julie A Harris, Lydia Ng, Brent Winslow, Nicholas Cain, Stefan Mihalas, Quanxin Wang, Chris Lau, Leonard Kuan, Alex M Henry, Marty T Mortrud, Benjamin Ouellette, Thuc N Nguyen, Staci A Sorensen, Clifford R Slaughterbeck, Wayne Wakeman, Yang Li, David Feng, Anh Ho, Eric Nicholas, Karla E Hirokawa, Phillip Bohn, Kevin M Joines, Hanchuan Peng, Michael J Hawrylycz, John W Phillips, John G Hohmann, Paul Wohnoutka, Charles R Gerfen, Christof Koch, Amy Bernard, Chinh Dang, Allan R Jones, and Hongkui Zeng, A mesoscale connectome of the mouse brain, Nature 508 (2014), 207–214.

76. James C Pang, Kevin M Aquino, Marianne Oldehinkel, Peter A Robinson, Ben D Fulcher, Michael Breakspear, and Alex Fornito, Geometric constraints on human brain function, Nature 618 (2023), 566–574.

77. Jonathan D Power, Kelly A Barnes, Abraham Z Snyder, Bradley L Schlaggar, and Steven E Petersen, Spurious but systematic correlations in functional connectivity mri networks arise from subject motion, NeuroImage 59 (2012), no. 3, 2142–2154.

78. Filippo Radicchi, José J Ramasco, and Santo Fortunato, Information filtering in complex weighted networks, Physical Review E 83 (2011), 046101.

79. Juan Luis Riquelme, Mike Hemberger, Gilles Laurent, and Julijana Gjorgjieva, Single spikes drive sequential propagation and routing of activity in a cortical network, eLife 12 (2023), e79928.

80. James A Roberts, Alistair Perry, Anton R Lord, Gloria Roberts, Philip B Mitchell, Robert E Smith, Fernando Calamante, and Michael Breakspear, The contribution of geometry to the human connectome, NeuroImage 124 (2016), 379–393.

81. Peter A Robinson, Physical brain connectomics, Physical Review E 99 (2019), 012421.

82. Frank Rosenblatt, The perceptron: a probabilistic model for information storage and organization in the brain., Psychological review 65 (1958), 386–408.

83. Mikail Rubinov and Olaf Sporns, Complex network measures of brain connectivity: uses and interpretations, NeuroImage 52 (2010), 1059–1069.

84. Mikail Rubinov and Olaf Sporns, Weight-conserving characterization of complex functional brain networks, NeuroImage 56 (2011), 2068–2079.

85. ML Saggio, P Ritter, and VK Jirsa, Analytical operations relate structural and functional connectivity in the brain, PLoS One 11 (2016), e0157292.

86. Toshikazu Samura, Yuji Ikegaya, and Yasuomi D Sato, A neural network model of reliably optimized spike transmission, Cognitive Neurodynamics 9 (2015), 265–277.

87. Louis K Scheffer, C Shan Xu, Michal Januszewski, Zhiyuan Lu, Shin-ya Takemura, Kenneth J Hayworth, Gary B Huang, Kazunori Shinomiya, Jeremy Maitlin-Shepard, Stuart Berg, Jody Clements, Philip M Hubbard, William T Katz, Lowell Umayam, Ting Zhao, David Ackerman, Tim Blakely, John Bogovic, Tom Dolafi, Dagmar Kainmueller, Takashi Kawase, Khaled A Khairy, Laramie Leavitt, Peter H Li, Larry Lindsey, Nicole Neubarth, Donald J Olbris, Hideo Otsuna, Eric T Trautman, Masayoshi Ito, Alexander S Bates, Jens Goldammer, Tanya Wolff, Robert Svirskas, Philipp Schlegel, Erika Neace, Christopher J Knecht, Chelsea X Alvarado, Dennis A Bailey, Samantha Ballinger, Jolanta A Borycz, Brandon S Canino, Natasha Cheatham, Michael Cook, Marisa Dreher, Octave Duclos, Bryon Eubanks, Kelli Fairbanks, Samantha Finley, Nora Forknall, Audrey Francis, Gary Patrick Hopkins, Emily M Joyce, SungJin Kim, Nicole A Kirk, Julie Kovalyak, Shirley A Lauchie, Alanna Lohff, Charli Maldonado, Emily A Manley, Sari McLin, Caroline Mooney, Miatta Ndama, Omotara Ogundeyi, Nneoma Okeoma, Christopher Ordish, Nicholas Padilla, Christopher M Patrick, Tyler Paterson, Elliott E Phillips, Emily M Phillips, Neha Rampally, Caitlin Ribeiro, Madelaine K Robertson, Jon Thomson Rymer, Sean M Ryan, Megan Sammons, Anne K Scott, Ashley L Scott, Aya Shinomiya, Claire Smith, Kelsey Smith, Natalie L Smith, Margaret A Sobeski, Alia Suleiman, Jackie Swift, Satoko Takemura, Iris Talebi, Dorota Tarnogorska, Emily Tenshaw, Temour Tokhi, John J Walsh, Tansy Yang, Jane Anne Horne, Feng Li, Ruchi Parekh, Patricia K Rivlin, Vivek Jayaraman, Marta Costa, Gregory SXE Jefferis, Kei Ito, Stephan Saalfeld, Reed George, Ian A Meinertzhagen, Gerald M Rubin, Harald F Hess, Viren Jain, and Stephen M Plaza, A connectome and analysis of the adult Drosophila central brain, eLife 9 (2020), e57443.

88. Caio Seguin, Martijn P van den Heuvel, and Andrew Zalesky, Navigation of brain networks, Proceedings of the National Academy of Sciences of the U.S.A. 115 (2018), 6297–6302.

89. M Ángeles Serrano, Marián Boguná, and Alessandro Vespignani, Extracting the multiscale backbone of complex weighted networks, Proceedings of the National Academy of Sciences of the U.S.A. 106 (2009), 6483–6488.

90. Taylor Simons and Dah-Jye Lee, A review of binarized neural networks, Electronics 8 (2019).

91. Olaf Sporns, Contributions and challenges for network models in cognitive neuroscience, Nature Neuroscience 17 (2014), 652–660.

92. Milou Straathof, Michel RT Sinke, Rick M Dijkhuizen, and Willem M Otte, A systematic review on the quantitative relationship between structural and functional network connectivity strength in mammalian brains, Journal of Cerebral Blood Flow and Metabolism 39 (2019), 189–209.

93. Laura E Súarez, Ross D Markello, Richard F Betzel, and Bratislav Misic, Linking structure and function in macroscale brain networks, Trends in Cognitive Sciences 24 (2020), 302–315.

94. Enzo Tagliazucchi, Pablo Balenzuela, Daniel Fraiman, and Dante R Chialvo, Criticality in large-scale brain fmri dynamics unveiled by a novel point process analysis, Frontiers in Physiology 3 (2012).

95. Naoya Takahashi, Takuya Sasaki, Wataru Matsumoto, Norio Matsuki, and Yuji Ikegaya, Circuit topology for synchronizing neurons in spontaneously active networks, Proceedings of the National Academy of Sciences of the U.S.A. 107 (2010), 10244–10249.

96. Andrew C Thomas and Joseph K Blitzstein, Valued ties tell fewer lies: Why not to dichotomize network edges with thresholds, (2011).

97. Cibu Thomas, Frank Ye, Okan Irfanoglu, Pooja Modi, Kadharbatcha Saleem, David Leopold, and Carlo Pierpaoli, Anatomical accuracy of brain connections derived from diffusion MRI tractography is inherently limited, Proceedings of the National Academy of Sciences of the U.S.A. 111 (2014), 16574–16579.

98. Anne E Urai, Brent Doiron, Andrew M Leifer, and Anne K Churchland, Large-scale neural recordings call for new insights to link brain and behavior, Nature Neuroscience (2022), 1–9.

99. Yezhou Wang, Jessica Royer, Bo-yong Park, Reinder Vos de Wael, Sara Larivìere, Shahin Tavakol, Raul Rodriguez-Cruces, Casey Paquola, Seok-Jun Hong, Daniel S Margulies, Jonathan Smallwood, Sofie L Valk, Alan C Evans, and Boris C Bernhardt, Long-range functional connections mirror and link microarchitectural and cognitive hierarchies in the human brain, Cerebral Cortex 33 (2023), 1782–1798.

100. Yu Wang, Eshwar Ghumare, Rik Vandenberghe, and Patrick Dupont, Comparison of different generalizations of clustering coefficient and local efficiency for weighted undirected graphs, Neural Computation 29 (2017), no. 2, 313–331.

101. Van J Wedeen, Ruopeng P Wang, Jeremy D Schmahmann, Thomas Benner, Wen Yih Isaac Tseng, George Dai, Deepak N Pandya, Patric Hagmann, Helen D’Arceuil, and Alexander J de Crespigny, Diffusion spectrum magnetic resonance imaging (DSI) tractography of crossing fibers, NeuroImage 41 (2008), 1267–1277.

102. Arthur T Winfree, The geometry of biological time, 2 ed., Springer, 2001.

103. Xiaoran Yan, Lucas GS Jeub, Alessandro Flammini, Filippo Radicchi, and Santo Fortunato, Weight thresholding on complex networks, Physical Review E 98 (2018), 042304.

104. Lina Yassin, Brett L Benedetti, Jean-Sébastien Jouhanneau, Jing A Wen, James F.A Poulet, and Alison L Barth, An embedded subnetwork of highly active neurons in the neocortex, Neuron 68 (2010), 1043–1050.

105. Andrew Zalesky, Alex Fornito, Luca Cocchi, Leonardo L Gollo, Martijn P van den Heuvel, and Michael Breakspear, Connectome sensitivity or specificity: which is more important?, NeuroImage 142 (2016), 407–420.

106. Changsong Zhou, Lucia Zemanová, Gorka Zamora, Claus C. Hilgetag, and Jürgen Kurths, Hierarchical organization unveiled by functional connectivity in complex brain networks, Physical Review Letters 97 (2006), 238103.

